# The PAXFOXO1s trigger fast trans-differentiation of chick embryonic neural cells into alveolar rhabdomyosarcoma with tissue invasive properties limited by S phase entry inhibition

**DOI:** 10.1101/2020.05.18.097618

**Authors:** Gloria Gonzalez Curto, Audrey Der Vartanian, Youcef Frarma, Line Manceau, Lorenzo Baldi, Selene Prisco, Nabila Elarouci, Frédéric Causeret, Muriel Rigolet, Frédéric Aurade, Aurélien De Reynies, Vincent Contremoulins, Frédéric Relaix, Orestis Faklaris, James Briscoe, Pascale Gilardi-Hebenstreit, Vanessa Ribes

## Abstract

The chromosome translocations generating PAX3FOXO1 and PAX7FOXO1 chimeric proteins are the primary hallmarks of the paediatric cancer, Alveolar Rhabdomyosarcoma (ARMS). Despite the ability of these transcription factors to remodel chromatin landscapes and promote the expression of tumour driver genes, they only inefficiently promote malignant transformation *in vivo*. The reason for this is unclear. To address this, we developed an *in ovo* model to follow the response of spinal cord progenitors to PAXFOXO1s. Our data demonstrate that PAXFOXO1s, but not wild-type PAX3 and PAX7, trigger the trans-differentiation of neural cells into ARMS-like cells with myogenic characteristics. In parallel expression of PAXFOXO1s remodels the neural pseudo-stratified epithelium into a cohesive mesenchyme capable of tissue invasion. Surprisingly, gain for PAXFOXO1s, as for wild-type PAX3/7, reduces the levels of CDK-CYCLIN activity and arrests cells in G1. Introduction of CYCLIN D1 or MYCN overcomes PAXFOXO1s mediated cell cycle inhibition and promotes tumour growth. Together, our findings reveal a mechanism underpinning the apparent limited oncogenicity of PAXFOXO1 fusion transcription factors and support a neural origin for ARMS.

## Introduction

Transcriptomic landscape remodelling represents a hallmark of tumourigenesis [1]. This is often achieved by perturbing the activity of powerful transcriptional modulators, such as master transcription factors (TFs). Understanding how the activity of these factors lead to a pathogenic transformation of cells represents a key challenge in cancer research, so is the development of *in vivo* and more physiological model systems to address this question [1,2].

Two related oncogenic TFs, PAX3FOXO1 and PAX7FOXO1, are associated with the emergence and development of the paediatric solid tumours, alveolar rhabdomyosarcoma (ARMS) [3]. Primary lesions in ARMS patients are mostly found in limb extremities or the trunk region. These comprise aggregates of round cells usually delineated by fibrous septa that express, as for other RMS subtypes, undifferentiated embryonic muscle cells markers. Along with these primary lesions, almost half of ARMS patients carry detectable metastases in the lung or bone marrow at the time of diagnosis. The occurrence of these metastases, together with cancer resistance and emergence of secondary disease are to blame for a poor cure rate of ARMS patients [4].

The in-frame pathognomonic chromosomal translocations, t(2;13)(q35;q14) or t(1;13) (p36;q14) fuse the 5’ end of the *PAX3* or *PAX7* genes to the 3′ end of the *FOXO1* gene and lead to the mis-expression of chimeric TFs made of the DNA binding domains of PAX3 or PAX7 TFs and the transactivation domain of FOXO1 [3]. Exome sequencing revealed that these translocations are the primary genetic lesions in more than 90% of ARMS cases [5,6]. Few somatic mutations are found in ARMS suggesting a relative fast development of the tumour after the translocations [6]. Furthermore, recurrent gross genetic aberrations, including whole genome duplication, unbalanced chromosomal copy gain, focal amplifications (12q13-q14 amplicon), or loss of heterozygosity notably on 11p15.5 locus presented by ARMS cells [5,6] suggest a tumorigenic transformation associated with chromothripsis [7]. The relative contribution of PAXFOXO1s and of these gross genetic aberrations during the transformation of healthy cells into ARMS cells is still debated.

A large body of work, mainly focused on PAX3FOXO1 and aimed at identifying and functionally characterizing PAXFOXO1’s target genes, argues the cell fate change characteristic of ARMS is driven by PAXFOXO1s [8,9]. This is hypothesised to arise from PAXFOXO1’s strong transcriptional transactivation potential, which surpasses that of normal PAX3 and PAX7 [10–13]. PAX3FOXO1 binds to non-coding *cis*-regulatory genomic modules (CRMs), remodelling chromatin and enhancing transcriptional activity [11,12]. These CRMs regulate the expression of genes associated with at least 3 traits deleterious for patients’ health [8,9,11,12,14]. First, several of the target genes encode cell surface proteins required for ARMS cells migration [15–19]. Second, ARMS cells harbour undifferentiated muscle cell determinants, which in presence of PAX3FOXO1 can no longer promote muscle terminal differentiation [20,21]. Third, PAXFOXO1s perturb the core cell cycle machinery [8,9]. Cross-interactions between PAX3FOXO1 with the anti-apoptotic gene *BCL-XL* or the senescent factor p16^INK4A^ promote cell survival [22–24], while PAX3FOXO1 activity during G2/M and G1/S checkpoints facilitates the amplification of transformed cells [25–27].

Despite the apparently powerful activity of PAXFOXO1s, data from animal models have led to the conclusion that the fusion proteins are not efficient in triggering ARMS formation and spreading [24,28–31]. In excess of 60 days are required for grafted PAX3FOXO1 expressing human myoblasts or mesenchymal stem cells to produce significant ARMS like growths in mice. This contrasts with the 15 days required for patient derived ARMS cells [30–32]. Similarly, driving PAX3FOXO1 expression in the muscle embryonic cells from the murine *Pax3* locus induces tumour mass with a reported frequency of 1 in 228 [13]. Importantly, these *in vivo* approaches have revealed several parameters enhancing PAXFOXO1 proteins oncogenicity. The random insertion of transgenes in zebrafish indicated that neural derived tissues are more prone than mesodermal derived tissues to produce tumours when exposed to PAX3FOXO1, highlighting the differential response of distinct cell lineages [29]. In addition, zebrafish and mouse models both indicate that a minimal level of PAX3FOXO1 needs to be reached to observe tumourigenesis [28,29,32]. Finally, complementing PAXFOXO1s expression with genetic aberrations promoting cell cycle progression markedly accelerated and increased the frequency of tumour formation [28–34]. This was notably achieved by lowering the expression of p53 or the retinoblastoma protein, RB1; or conversely by ectopically elevating MYCN expression or RAS activity.

To investigate the molecular mechanisms of oncogenicity in ARMS we characterised the initial cellular and molecular steps associated with the transformation of cells expressing PAX3FOXO1 and PAX7FOXO1. The growing evidence for an embryonic origin of paediatric cancers [35], the identification of ARMS growths in neural tube derived tissues [36,37], the recurrent presence of embryonic neural lineage determinants in ARMS cells [9], and the recent use of chick embryos to study cancer cells migration and invasion [38,39] led us to develop the embryonic chick neural tube as a model system. We demonstrate that PAXFOXO1s repress the molecular hallmarks of neural tube progenitors within 48 hours and impose a molecular signature reminiscent of that of ARMS cells. Concomitantly, PAXFOXO1s promote an epithelial-mesenchymal transition, conferring cells with the ability to invade the adjacent mesoderm in less than 72 hours. Moreover, PAXFOXO1s limit cell cycle progression, *via* a reduction of CDK-CYCLIN activity, which in turn can explain the limited oncogenicity of these fusion TFs.

## Results

### Chick neural cells lose their neurogenic potential upon PAXFOXO1 exposure

To investigate the transformation potential of PAXFOXO1 proteins, we set out to perform gain of function experiments *in vivo* using the neural tube of chick embryos. Hamburger and Hamilton (HH) stage 11 chick embryos were electroporated with a vector expressing either *PAX3FOXO1* or *PAX7FOXO1* together with a bi-cistronically encoded nuclear-targeted GFP and allowed to develop *in ovo* for up to 72h (Fig 1A). For comparison, electroporations with the wild-type versions of *Pax3* and *Pax7* or the empty *pCIG* vector were performed.

**Fig 1:**
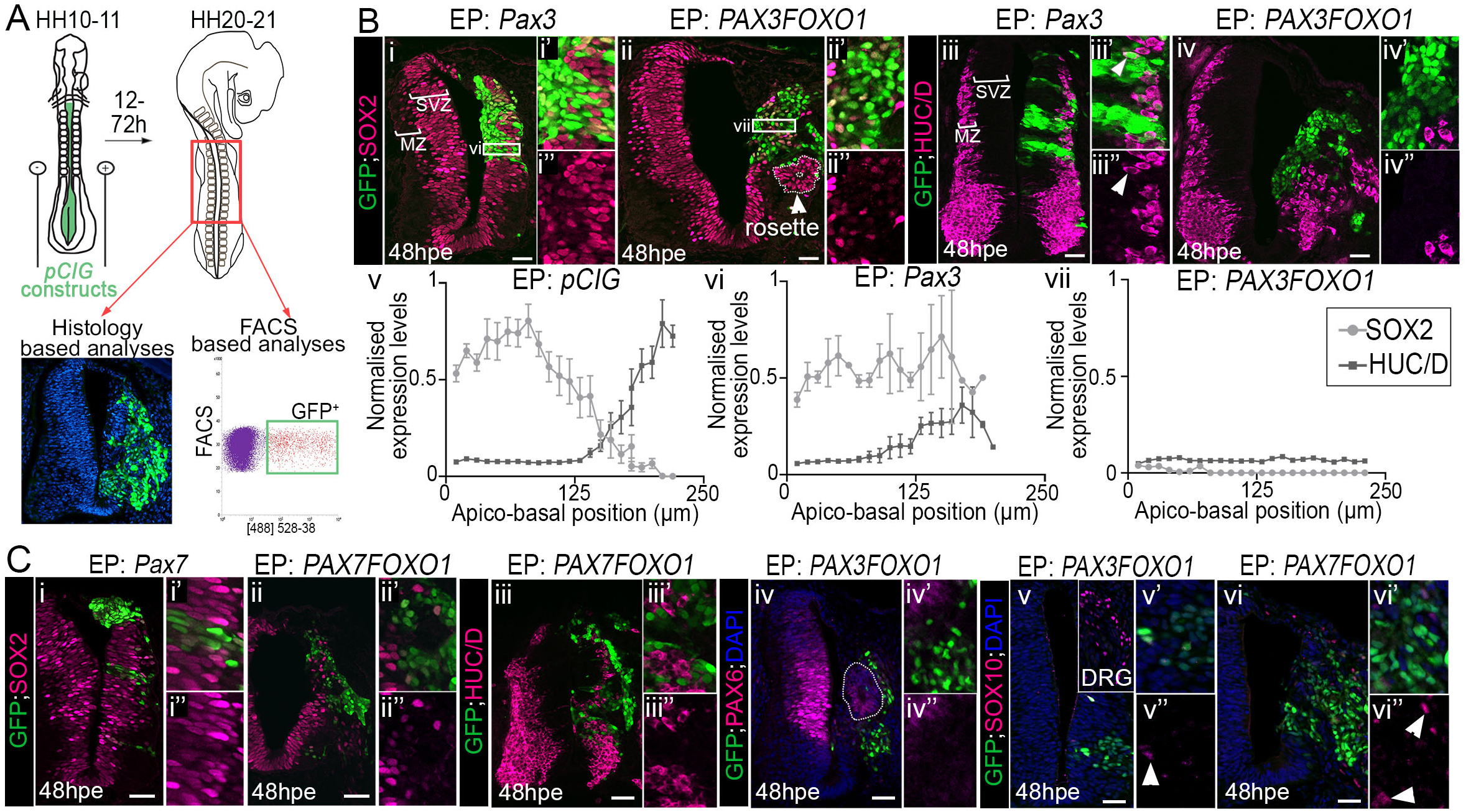
PAXFOXO1s switch off generic neurogenic marker expression in chick embryonic spinal cord. **(A)** Schematics showing HH10-11 chick embryos filled with *pCIG* based constructs before being electroporated. The cervical to thoracic region (red square) of the electroporated embryos were dissected 12 to 72 hours later to perform histology or FACS based analyses. **(B)(i-iv’’)** Immunodetection of GFP, SOX2 and HUC/D on transverse sections of chick embryos 48hpe with *Pax3* or *PAX3FOXO1.* MZ: Mantle Zone; SVZ: Subventricular Zone; Arrow in ii points at a rosette of SOX2^+^;GFP^-^ cells clustering apart from the SVZ. **(v-vii)** Quantification of SOX2 and HUC/D expression levels 48hpe in bins containing only GFP^+^ cells and positioned along the apico-basal axis and within the dorsal neural tube as described in materials & methods section (cf rectangles depicted in i and ii). **(C)** Immunodetection of GFP, SOX2, HUC/D, PAX6, SOX10 and DAPI staining on transverse sections of chick embryos at 48hpe with the indicated constructs. DRG: dorsal root ganglia. Arrows in v’,vi’’ point to rare SOX10^+^;GFP^+^ cells. **Generals**: scale bars: 50µm. x’ and x’’ panels are blown up on a subset of GFP^+^ cells present in the panel x.

We first characterised the molecular identity of electroporated cells by assaying the expression of generic neuronal markers (Fig 1B,C). At 48 hours post electroporation (hpe), the neural tube of chick embryos contained SOX2^+^ progenitors located closed to the ventricle and HUC/D^+^ neurons laterally in the mantle zone (brackets in Fig 1Bi,iii). PAX3FOXO1 and PAX7FOXO1 overexpression caused a marked reorganisation on both the ventricular and mantle regions of the neural tube (Fig 1Bii-ii’’,iv-iv’’,vii, Cii-vi’’). Strikingly, PAXFOXO1^+^ cells lacked both SOX2 and HUC/D. Hence, the fusion TFs appear sufficient to reprogram cells engaged into generic neurogenic program. This is further sustained by the strong PAX3FOXO1 mediated reduction in the expression levels of PAX6, another pan-neuronal progenitor marker (Fig 1Civ-iv’’). We also checked for the expression of SOX10, a marker of neural crest cells (NCC)[40]. These cells are specified from the dorsal most part of the neural tube, which they leave to colonize distant embryonic tissues, including the dorsal root ganglia (DRG). At 48hpe and on the un-electroporated side of the embryo, SOX10^+^ NCC were present in the skin and the DRG (inset in Fig 1Cv). On the electroporated side, only rare PAXFOXO1^+^ cells were also positive for SOX10 (arrows in Fig 1Cv-vi’’), ruling out the possibility of a switch of neural cells into NCC upon exposure to the fusion TFs.

By contrast to the fusion proteins, PAX3 or PAX7 overexpression did not affect the organisation of the neural tube and cells kept expressing neurogenic factors, such as SOX2 (Fig 1Bi-i’’,vi, Ci-i’). Yet, in some cells expressing high levels of the PAX3 or PAX7, SOX2 expression levels were reduced. This is consistent, with both PAX3 and PAX7 been present in progenitors of the dorsal half of the neural tube. More significantly, less HUC/D^+^ neurons were generated in the neural tube overexpressing *Pax3* (Fig 1Biii-iii”); a phenotype reminiscent to the forced expression of another PAX TF member, PAX6 and suggesting that the extinction of PAX is required in neural progenitors for their terminal differentiation [41].

### PAXFOXO1 TFs convert chick neural cells into ARMS like-cells

We next tested whether PAXFOXO1 expressing cells adopted the identity of alveolar rhabdomyosarcoma cells. To refine the list of genes that define this identity [9], we combined and re-analysed microarray-based tumour transcriptomes obtained from 133 PAXFOXO1 positive ARMS patients and 59 patients affected by other RMS subtypes (Fig 2A, S1 Table, S1A Fig)[42–46]. We identified 1194 genes enriched in ARMS biopsies; 40% of which were in the vicinity of previously identified PAX3FOXO1 bound *cis*-regulatory modules (CRM) [11,12](Fig 2B). This list of genes largely comprised previously identified PAX3FOXO1 dependent ARMS markers, such as *ALK, ARHGAP25*, or *FGFR4* [9,47]. Functional annotation of these genes indicated that they encode for developmental regulators of many embryonic lineages known to be dependent on PAX3 and PAX7 activities (Fig 2C, Table S2)[48], and not exclusively of the muscle lineage. ARMS signature was composed of TFs which are not normally co-expressed in vertebrate embryos. For instance, during early/late somitogenesis in the caudal part of amniotes, *LMO4*[49], *MEOX1*[50], *MYOD1*[51], *PITX2*[52] label embryonic muscle cells, *FOXF1*[53] is in the sclerotome and *PAX2*[54], *PRDM12*[55], *TFAP2α*[56] mark neurons of the peripheral and/or central nervous system. This suggests that ARMS cells are not simply undifferentiated muscle cells, but rather as cells with their own transcriptional status. To verify the presence of this combination of TFs in ARMS, we quantified their expression levels using either RT-qPCR or western blots across seven established human RMS cell lines, including 3 embryonal rhabdomyosarcoma (ERMS; RD, RDAbl, Rh36) and 4 PAX3FOXO1 positive ARMS cell lines (Rh30, SJRh30, Rh4, Rh5) (Fig 2D, S1B Fig). All markers assessed were present in ARMS cell lines, with expression levels varying from one cell line to another. Nevertheless, most of them displayed elevated levels in ARMS cells compared to ERMS cells. This combination of TFs represents, thus, a good hallmark of ARMS cell identity, an identity mixing determinants of distinct embryonic lineages.

**Fig 2:**
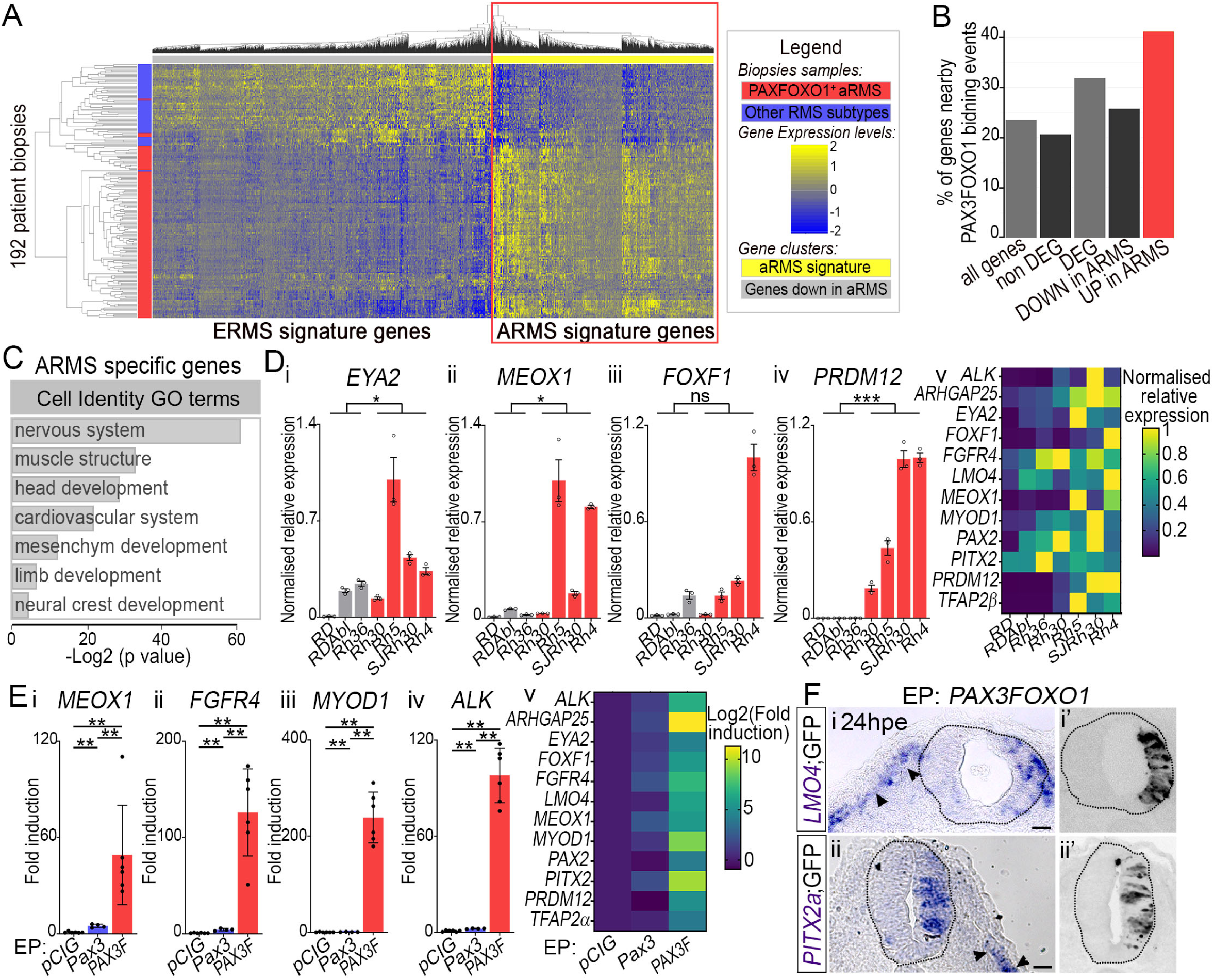
PAXFOXO1 TFs convert embryonic neural progenitors into cells harbouring ARMS molecular traits. **(A)** Heatmap of hierarchically clustered differentially expressed genes between PAXFOXO1 positive ARMS (red rectangles) and other RMS biopsies (blue rectangles). Fold changes across samples are colour-coded in blue (lower levels) to yellow (higher levels) (See also Method section and S1 Table). Genes upregulated in ARMS versus ERMS are squared in red and named ARMS signature genes. **(B)** Percentage of genes present nearby at least one known PAX3FOXO1 bound CRM [12] out of those present in our complete transcriptomic data set (all genes), non-differentially regulated between ARMS and other RMS (non DEG), the differentially expressed genes between ARMS and other RMS (DEG), downregulated in ARMS compared to other RMS (DOWN in ARMS), upregulated in ARMS compared to other RMS (UP in ARMS). **(C)** Gene ontology enrichment for biological processes terms linked to tissue specification applied to genes enriched in ARMS biopsies. **(D)** mRNA expression levels of ARMS signature genes nearby a PAX3FOXO1 binding event and expressed in various PAX3/7 dependent embryonic tissues assayed by RT-qPCR on the indicated ERMS and ARMS cell lines. Levels are relative to *TBP* transcripts and normalised to max value across all cell lines. **i-iv:** dots: value for a single RNA preparation; histograms: mean ± s.e.m.; n=3; **v:** heatmap displays mean value in each cell line. Normalised relative expression across samples are colour-coded in blue (lower levels) to yellow (higher levels). **(E)** mRNA expression levels of ARMS hallmark genes in GFP^+^ FACS sorted neural tube cells 48hpe with *pCIG, Pax3* and *PAX3FOXO1*. Levels are relative to *TBP* transcripts and normalised to *pCIG* samples mean level. **i-iv:** dots: value for a single RNA sample; histograms: mean ± s.e.m; n=6; **v:** heatmap exhibits mean value over 4 discrete FAC sorts. Fold induction across samples are colour-coded in blue (lower levels) to yellow (higher levels). **(F)(i, ii)** *LMO4* and *PITX2* detection via *in situ* hybridization on transverse sections of chick embryos 24hpe with *PAX3FOXO1* and (i’, ii’) immuno-detection of GFP on the adjacent section slide. Black arrows indicate normal localisation of cells expressing *LMO4* and *PITX2a*. n>6 embryos. **Generals:** p-values evaluated using two-way-ANOVA (D) or Mann-Whitney U test (E): *: p<0.05, **: p< 0.01, ***: p<0.001, ****: p<0.0001, ns: p>0.05; Scale bars: 50µm.

We next assessed the expression of these ARMS hallmark genes in *GFP, Pax3* or *PAX3FOXO1* electroporated chick neural cells (Fig 2E). For this, the neural tube of 48hpe embryos were dissected, dissociated and FACS purified (Fig 1A). RNA from 60 to 80k GFP positive cells were extracted and RT-qPCR used to assay specific cDNAs. The expression of all genes was significantly increased by PAX3FOXO1 and barely altered by PAX3 (Fig 2E). It is noteworthy that determinants of the muscle cells were as much induced as genes marking other cell types in embryos. *In situ* hybridization for *PITX2* and *LMO4* confirmed the ectopic induction of these genes by PAX3FOXO1 with levels of expression reminiscent of those found in somites derived structures (Fig 2F). Similarly, immunofluorescent staining for TFAP2α and PAX2, showed that these two proteins are induced in more than 70% of PAXFOXO1 electroporated cells (S1C Fig). Conversely, forced expression of PAX3 or PAX7 tented to reduce the number of TFAP2α^+^ and PAX2^+^ neurons generated. Altogether our data provides evidence that PAXFOXO1 factors have the ability to induce a molecular signature reminiscent of human ARMS cells in neural cells, a non-muscle lineage.

### PAXFOXO1s activate conserved ARMS associated enhancers in chick neural cells

The robustness of PAXFOXO1 mediated ARMS hallmark gene induction in neural cells could stem from the activation of conserved enhancers known to operate in ARMS cells [11,12]. To test this idea, we amplified enhancers found in the vicinity of the mouse *Met, Meox1, Myod1, Alk*, or human *CDH3* and *PRDM12* genes. We cloned these upstream of a minimal promoter and a reporter gene and co-electroporated them with either *Pax3, Pax7, PAX7FOXO1, PAX3FOXO1* or *pCIG* (Material and Methods). In the neural tube of embryos electroporated with the control vector the activity of these enhancers was barely detectable (Fig 3Ai,i’,iv,iv’), with the exception of one CRM near the *PRDM12* locus that had an endogenous activity in the intermediate-dorsal neural tube (Fig 3Bi,i’). For most CRM, the wild-type PAX had moderate effects on their activity (Fig 3Aii,ii’,v,v’,vii, Biii). In contrast, upon PAX3FOXO1 expression, all cloned enhancers, except the *CDH3* CRM, were active and promoted the expression of the reporter genes (Fig 3A-C). The magnitude of responses varied between enhancers and from cell to cell. Testing the effects of PAX7FOXO1 on the activity of *PRDM12*^*CRM*^ showed that the activity of this fusion protein was similar to PAX3FOXO1 (Fig 3Biii). Altogether these results support a model whereby the transformation of neural progenitors to an ARMS cell identity could be mediated by the co-option of conserved enhancer elements.

**Fig 3:**
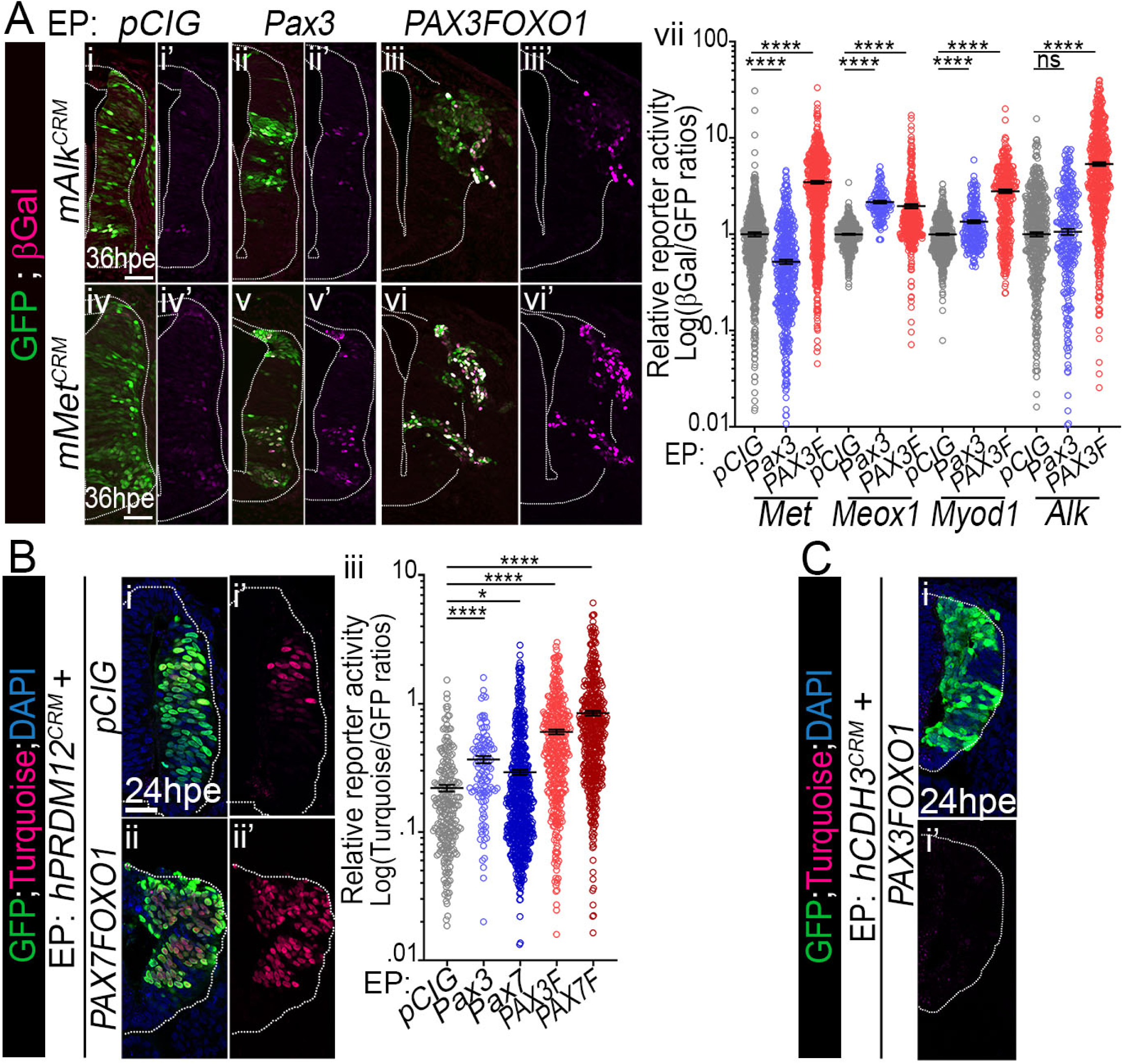
Activation of ARMS associated enhancers in chick neural cells by PAXFOXO1 TFs. **(A)** (**i-vi’**) Immunostaining for GFP and βGalactosidase (βGal) on transverse sections of chick embryos 36hpe with *pCIG, Pax3* or *PAX3FOXO1* and the indicated reporters for the mouse versions of *cis*-regulatory modules (CRM) bound by PAX3FOXO1 in ARMS cells [12]. (**vii**) Quantification of βGal levels normalised to that of GFP in cells electroporated with the indicated enhancer reporter constructs at 36hpe (dots: single cell values; bars: mean ± s.e.m.; n>4 embryos). **(B-C)** (**i-ii’**) Immunostaining for GFP, Turquoise direct fluorescence and DAPI staining on transverse sections of chick embryos 24hpe with *pCIG* or *PAX7FOXO1* or *PAX3FOXO1* and a reporter for human *PRDM12*^*CRM*^ and *CDH3*^*CRM*^. (**iii**) Quantification of Turquoise levels normalised to that of GFP in cells electroporated with *PRDM12*^*CRM*^ construct at 24hpe (dots: single cell values; bars: mean ± s.e.m.; n>4 embryos). **Generals**: scale bars: 50µm; Mann-Whitney U test p-value: *: p< 0.05, ****: p< 0.0001, ns: p> 0.05.

### PAXFOXO1 TFs promote epithelial-mesenchymal transition, cell migration and tissue invasion

Paralleling PAXFOXO1 mediated cell fate changes, drastic rearrangement of the pseudo-stratified neuro-epithelium occurred (Fig 4Aiii, iii’, S2Aii,ii’ Fig). PAXFOXO1^+^ cells adopted a rounded shape, were unevenly distributed within the tissue, yet grouped together. Some cells had delaminated either inside the neural tube canal or within the adjacent mesodermal tissue. In addition, neighbouring unelectroporated cells clustered together to form SOX2^+^ rosettes ectopically positioned within the “mantle zone”, supporting a sorting of PAXFOXO1^+^ cells from the non-electroporated ones (arrows in Fig 1Bii, 4Aiii,iii’, S2Aii,ii’ Fig). In contrast, *pCIG* expressing, PAX3^+^ and PAX7^+^ cells were aligned, elongated and confined to the neural tube (Fig 4Aii-ii’, S2Ai,i’ Fig). In addition, PAX3^+^ and PAX7^+^ cells electroporation resulted in a thinner neuro-epithelium than seen in *pCIG* samples (Fig 4Ai-ii’, S2Ai,i’ Fig).

**Fig 4:**
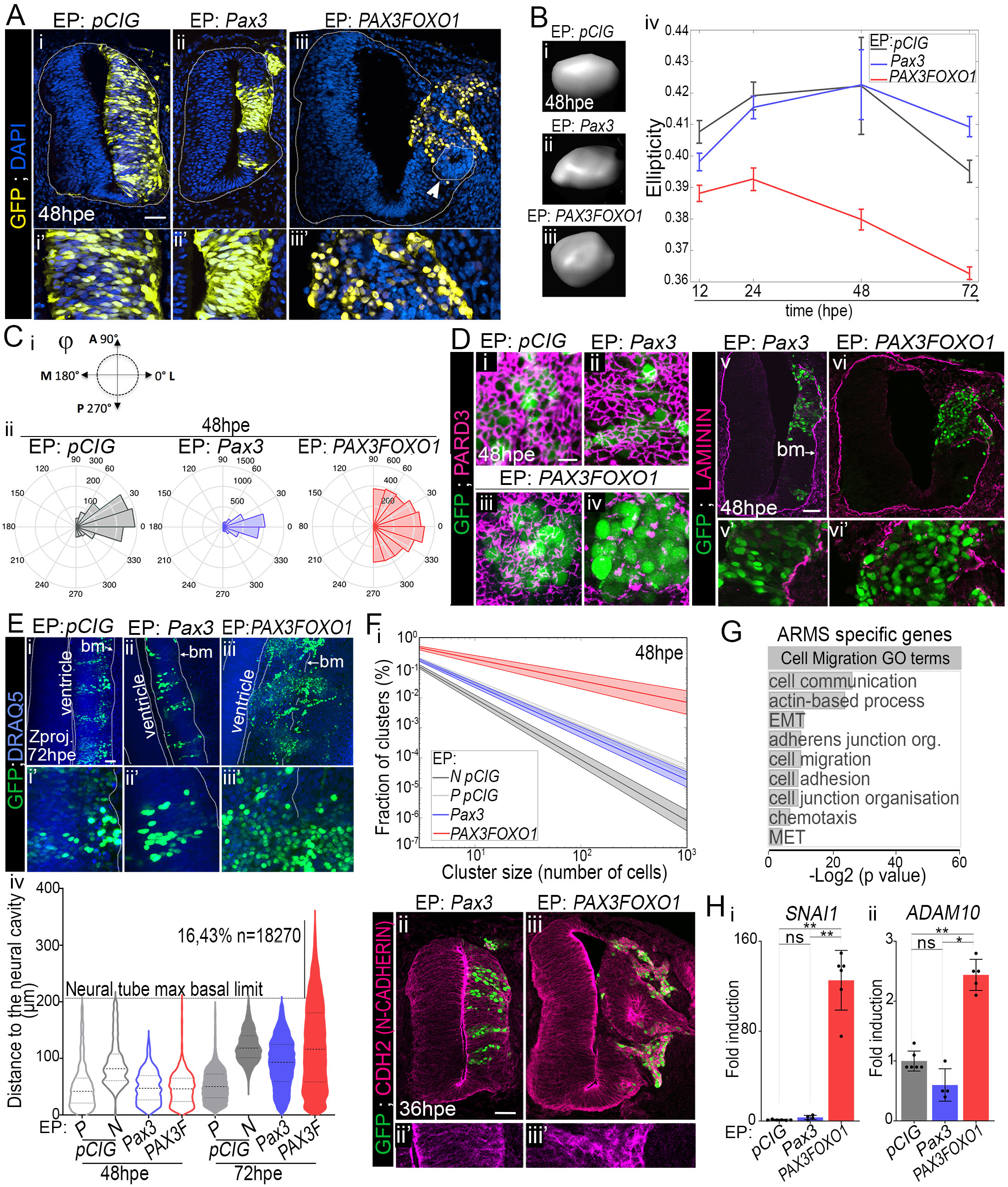
PAXFOXO1 TFs transform neural epithelial cells into a cohesive mesenchyme capable of migration. **(A) (i-iii’)** Immunodetection of GFP and DAPI stainings on transverse section of chick embryos 48hpe with the indicated plasmids. **(B) (i-iii)** Representative 3D shape of GFP^+^ nuclei segmented from scanned whole embryos 48hpe with the indicated plasmids. **(iv)** Temporal dynamics of the ellipticity of nuclei measured from the segmentation of GFP^+^ nuclei (as shown in i-iii) in whole-mount embryos (mean ± s.d., n>6 embryos). **(C) (i)** φ possible values and major axes of chick embryos it refers to. A: anterior; P: posterior; M: medial; L: lateral **(ii)** Distribution of *φ* in embryos electroporated with the indicated constructs at 48hpe. **(D) (i-iv)** Apical confocal views in open-booked preparation of spinal cords of embryos 48hpe with the indicated constructs and immunolabelled with antibodies against GFP and PARD3. Variations in the phenotype are observed in presence of PAX3FOXO1. **(iii)** represents 2 of 8 cases analysed, **(iv)** the rest of cases. (**v-vi’**) Immunodetection of GFP and LAMININ on transverse sections of chick embryos 48hpe with the indicated plasmids. **(E)** (**i-iii**) Z-projections of whole mount embryos immuno-labelled for GFP and stained with DRAQ5. Dotted lines delineate either the neural tube cavity of the neural tube/mesoderm border. **(iv)** Quantification of the distance of each GFP^+^ nuclei from the apical surface at 48hpe and 72hpe with the indicated plasmids (Violin plots) P: progenitors and N: neurons. **(F) (i)** Exponential fit of the % of cell clusters as the function of cluster size at 48hpe in discrete sample types. **(ii**,**iii)** Immunodetection of GFP and N-CADHERIN on transverse sections of chick embryos 48hpe with the indicated plasmids. **(G)** Gene ontology enrichment for biological processes terms linked cell migration and adhesion applied to genes enriched in ARMS biopsies. EMT: epithelial to mesenchymal transition; MET: mesenchymal to epithelial transition. **(H) (i-ii)** Levels of mRNA expression of the indicated ARMS signature gene assayed by RT-qPCR on GFP^+^ FACS sorted neural tube cells 48hpe with *pCIG, Pax3* and *PAX3FOXO1*. Levels are relative to *TBP* transcripts and normalised to *pCIG* samples mean level (dots: value for a single RNA sample; histograms: mean ± s.d.; Mann-Whitney U test p-value: *: p<0.05, **: p<0.01, ns: p>0.05). **Generals:** Scale bars: 50µm, but in D: 10µm. Bottom x’ panels are blown up of the x panels.

To validate these observations, we quantified several parameters in whole embryos stained with the DNA dye DRAQ5 and GFP and documented the distribution of several key markers of the epithelial state (S2B Fig). The tumour cell modes of migration are tightly connected to cell shape (e.g. [57]). Hence, we started by evaluating that of GFP^+^ cells by measuring the ellipticity of their nuclei segmented from 3D images (Fig 4B). This parameter reflects the degree of divergence from a sphere. It fluctuated between 0.4 and 0.42 for *pCIG* and *Pax3* elongated nuclei. The ellipticity of PAX3FOXO1^+^ cells was substantially smaller; with time this difference was accentuated. PAX3FOXO1 cells adopt a rounded shape likely adapted to tissue exploration [57].

We then monitored the orientation of the major axis of the ellipsoid fit of GFP^+^ cells using polar coordinates, a parameter indicative of cell arrangement within the tissue (Fig 4C, S2C Fig). The polar angle *θ* gave the deviation of the nuclei major axis to the tissue dorso-ventral axis, while the azimuthal angle *φ* informed on its orientation within the lateral-medial and posterior-anterior tissue plane (S2Ci,ii Fig). In controls and *Pax3* samples, the distribution of *θ* and *φ* barely fluctuated over time (Fig 4Cii, S2Ciii Fig). *θ* was centred on 90°C, *φ* on 0°C, consistent with nuclei paralleling the medial-lateral axis of the embryos and apico-basal attachments of cells. In contrast, in *PAX3FOXO1* samples *θ* and *φ* values displayed a wide distribution (Fig 4Cii, S2Ciii Fig), ranging for instance for *φ* between −90° and + 90°. This variability was set as soon as 12hpe and increased with time (S2Ciii Fig). Hence, PAX3FOXO1 is able to randomize the nuclei orientation within the spinal tissue from 12hpe onwards.

Alterations in the shape and orientation of the nuclei by PAX3FOXO1 led us to assess the apical-basal attachment of cells (Fig 4D, S2D Fig)[58]. We monitored the distribution of the apical determinant PARD3. In open book preparations of 48hpe whole spinal cord, PARD3 labelling revealed a honeycomb-like network at the apical surface (Fig 4Di). This network remained intact upon gain for PAX3 although cells harboured less cell-cell contacts (Fig 4Dii). In contrast, PAX3FOXO1 completely disassembled this network (Fig 4Diii,iv). Barely detectable at 24hpe, this phenotype became evident by 48hpe (S2Di-iv’’ Fig). The loss of apical polarity was confirmed by looking at the apically located activated form of βCATENIN and ARL13B^+^ primary cilia (S2Di-v’’ Fig). We next looked at the distribution of the focal adhesion anchor β1 INTEGRIN, which accumulates within the basal regions of control cells (S2Dvi-vi’’ Fig). The expression of this protein was homogenous throughout PAX3FOXO1^+^ cells (Fig S2Dvi,vi’’). Hence, upon PAXFOXO1 expression, neural progenitors lose the polarized distribution of cell-cell and cell-matrix attachments that become distributed evenly throughout their cell membrane.

Because cell polarity is influenced by the extra cellular matrix (ECM)[58], we investigate the distribution of LAMININ (Fig 4Dv-vi’). This key scaffold component of the basal lamina separates the neural tube from the adjacent mesoderm, as shown in *Pax3* samples at 48hpe (Fig 4Dv,v’). In the presence PAX3FOXO1, the basal lamina broke down (Fig 4Dvi,vi’; S2D Fig); a phenotype progressively appearing from 24hpe onwards. This indicated that PAXFOXO1s provide cells with the ability to dismantle tissue barriers. We next tested whether PAXFOXO1^+^ cells were capable of migration, by measuring the distances between the centre of electroporated nuclei and the apical surface of the neural tube in the 3 dimensions of embryos (Fig 4Ei-iv). In *pCIG* samples, the arrangement of progenitors and neurons nuclei differed and allowed to distinguish these two types of cells. This could not be done in *Pax3* samples, probably because too few neurons were produced (Fig 1B) and in PAX3FOXO1 due to the global transformation of cells (Fig 1B-C). PAX3 nuclei remained in the neural tube at both 48 and 72hpe. Similarly, up to 48hpe, PAX3FOXO1^+^ cells were contained within the neural tube. 24 hours later, a fraction of these cells (more than 15%) had migrated outside the neural tube and were present within the adjacent tissues. This was also confirmed for PAX7FOXO1 using transverse section analysis (S2E Fig), fully validating the idea that PAXFOXO1 fusion proteins provide cells with invasive properties.

To investigate whether cells clustered together, we first measured the distance between nearest nuclei from which we evaluated the number of cells belonging to the same cluster (Fig 4Fi, see Methods). In control embryos, electroporated neuro-progenitors were more clustered together than neurons, which is in agreement with the delamination and various migration paths taken by neuronal subpopulations. PAX3 electroporated cells behave similarly to control neural progenitors (Fig 1B). By contrast, in PAX3FOXO1 electroporated neural tubes, we identified more cells close to each other and bigger groups of cells than in control, supporting the idea that it favours the clustering of cells. In agreement with this, PAX3FOXO1^+^ cells expressed high levels of N-CADHERIN (CDH2), which was homogenously distributed throughout the cells (Fig 4Fii-iii’).

Taken together, these data indicate that PAXFOXO1 fusion genes not only trigger acquisition of ARMS identity markers but also provide cells with the ability to invade tissue. These phenotypes are likely to be directly regulated by PAXFOXO1s, as suggested by the great number of PAXFOXO1 targets in ARMS cells encoding for tissues remodellers and cell migration regulators (Fig 4G, S2F Fig). We notably confirmed that the master epithelial-mesenchymal transition driver *SNAI1* and the ECM remodeller *ADAM10* genes displayed elevated levels in presence of PAX3FOXO1 compared to control and PAX3^+^ chick neural cells (Fig 4H).

### PAXFOXO1 TFs hold cells in G1 by decreasing CDK-CYCLIN activity

We next assessed the impact of PAXFOXO1 proteins on other hallmarks of cancer cells, notably those related to their cycling behaviour [14](Fig 5, S3 Fig).

**Fig 5:**
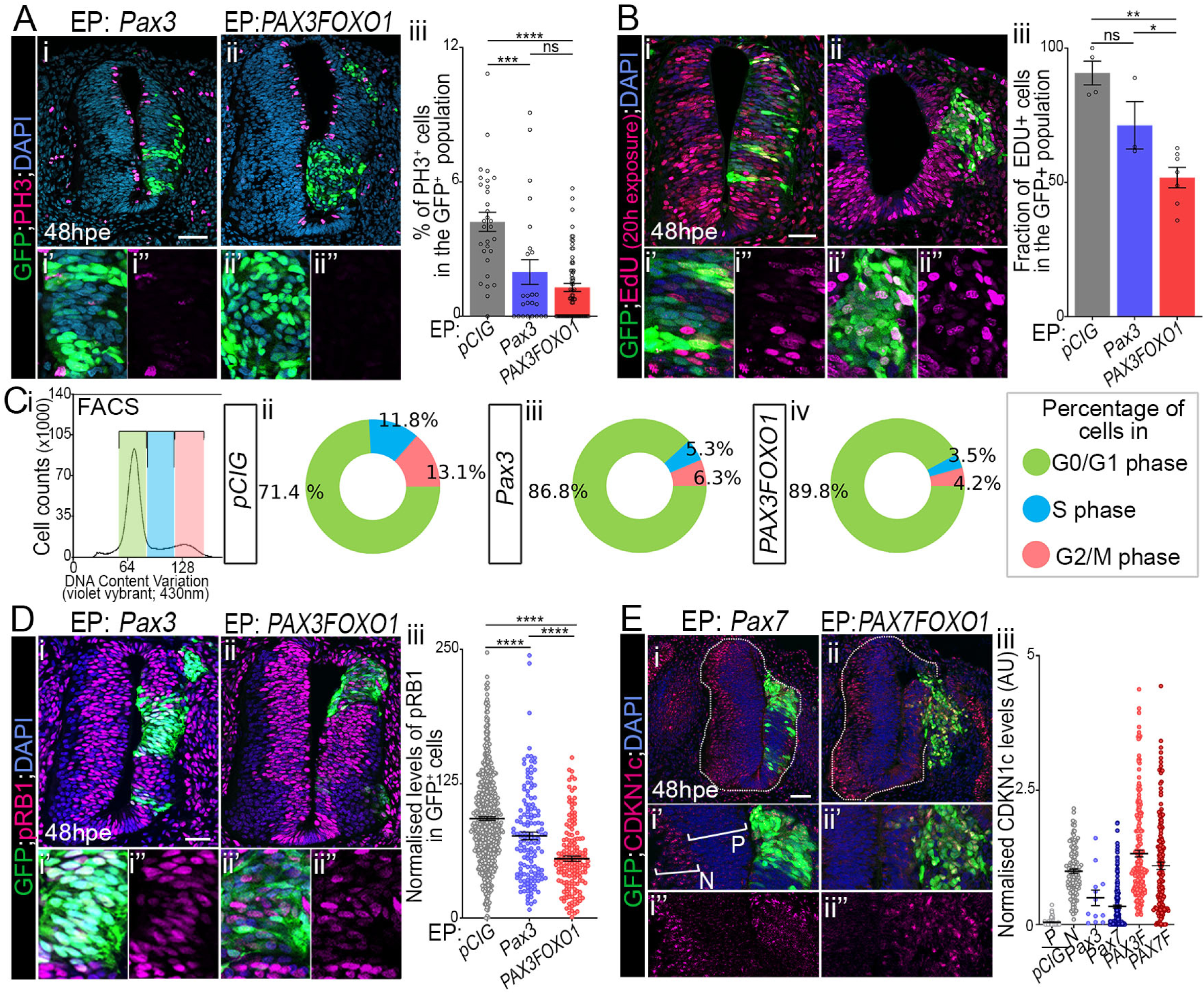
Pax3/7 and PAXFOXO1 TFs limit the entry of cells into S phase. **(A) (i-ii’)** Immunodetection of GFP, phospho-histone H3 (PH3) and DAPI staining on transverse sections of chick embryos 48hpe with the indicated plasmids. **(iii)** Quantification of the number of PH3^+^ cells in the GFP^+^ population in embryos expressing the indicated plasmids (dots: values on one section, n>8 embryos; histograms: mean ± s.e.m.). **(B) (i-ii’)** Immunodetection of GFP, EdU and DAPI staining on transverse sections of chick embryos 48hpe with the indicated plasmids and soaked with EDU 20h before harvest. **(iii)** Quantification of the number of EdU^+^ cells in the GFP^+^ population in embryos expressing the indicated plasmids (dots: values on one section, n=3 embryos; histograms: mean ± s.e.m.). **(C)(i)** FACS plots showing DNA content distribution of GFP^+^ chick neural cells stained with violet vybrant dye and cell cycle phases gating (green: G0/G1 phase, blue: S phase and pink: G2/M phase). **(ii-iv)** Percentage of cells in the indicated cell cycle phase at 48hpe established as in i (mean over three experiments, for individual values see Fig S3Ei-iii). **(D) (i-ii’)** Immunodetection of GFP, phosphorylated form of RB1 (pRB1) and DAPI staining on transverse sections of chick embryos 48hpe with the indicated plasmids. **(iii)** Quantification of pRB1 in the GFP^+^ population in embryos expressing the indicated plasmids (dots: values in arbitrary unit (AU) on one section, n> 8 embryos; bars: mean ± s.e.m). **(E) (i-ii’)** Immunodetection of GFP, CDKN1c and DAPI staining on transverse sections of chick embryos 48hpe with the indicated plasmids. P: progenitors; N: neurons. **(iii)** Quantification of the CDKN1c levels in GFP^+^ cells in embryos expressing the indicated plasmids P: progenitors; N: neurons. (dots: values on one section, n=3 embryos; bars: mean ± s.e.m.). **Generals:** Bottom panels are blown up the squares shown in upper ones; Scale bars: 50µm; Mann-Whitney U test p-value: *: p< 0.05, **: p< 0.01, ***: p< 0.001, ****: p< 0.0001, ns: p>0.05.

To assess the proliferative state of cells, we marked mitotic cells using an antibody against the phosphorylated form of histone H3 (PH3) (Fig 5A, S3A Fig). This indicated PAXFOXO1^+^ cells displayed a lower rate of mitosis than control cells at both 24 and 48hpe. Such a reduction in the number of PH3^+^ cells was seen also in PAX3/7^+^ cells, albeit to a lesser extent. These results suggest that either PAXFOXO1^+^ cells were blocked in a cell cycle phase or had a longer cell cycle(s). We first evaluated if the fusion proteins specifically induce cell death by marking activated CASPASE3^+^ apoptotic cells (S3B Fig). Upon overexpression of any of the *Pax3/7* variants, a too low proportion of cells (about 2%) were undergoing cell death at 48hpe to account for the mitotic rate decrease. We next traced cells undergoing DNA synthesis by treating embryos with EdU for 20h before harvesting. Nearly all control cells were positive for EdU, while only half of PAX3FOXO1^+^ cells and 75% of PAX3^+^ cells incorporated the thymidine analogue (Fig 5B). Confirming this compromised entry into replication, the expression levels of minichromosome maintenance 2 (MCM2), a protein of the pre-replicative complex was significantly downregulated in cells expressing PAX3/7 and even more in PAXFOXO1 cells (S3D Fig). FACS analyses, indicated that in presence of PAX3 and PAX3FOXO1 a larger proportion of cells were in the G1 phase (Fig 5C). Taken together, these results support the idea that the gain for PAX3/7 or PAX3/7FOXO1 arrests cells in G1 phase. Similar experiments performed in human fibroblasts indicated that this cell type was also arrested in G1 upon gain for PAX3FOXO1 (S3E Fig), supporting the idea that PAXFOXO1 mediated cell cycle blockade is not inherent to our chick system.

Finally, the phosphorylation of the retinoblastoma-associated RB1 protein being one of the hallmarks of the CDK-CYCLIN activity leading to the entry in S phase, we assayed its status. All PAX3/7 variants decreased phospho-Rb1 levels; the fusion proteins to a greater extent than wild-type PAX (Fig 5D, S3C Fig). This is phenotype is not linked to a decrease in the transcription of *RB1, CDK2, CDK6* and *CCND1* (S3F Fig). Instead, we identified that amongst the CIP/KIP CDK inhibitors, CDKN1c (P57Kip2) was upregulated (Fig 5E), a cue potentially explaining the PAXFOXO1 mediated decrease in CDK-CYCLIN activity.

### PAXFOXO1 mediated cell cycle inhibition is overcome by CCND1 or MYCN

We then wanted to test whether PAXFOXO1-transformed cells could re-enter cell cycle. For this, we first decided to reactivate CDK-CYCLIN activity in PAX3FOXO1 expressing cells, by forced expression of CYCLIN D1, CCND1 [59]. In presence of this cyclin subtype, PAX3 and PAX3FOXO1 positive cells displayed a mitotic rate, revealed by quantifying PH3^+^ cells, similar to that of *pCIG* control embryos at both 24 and 48hpe (Fig 6A). Accordingly, the gain for CYCLIN D1 allowed PAX3FOXO1^+^ cells to incorporate EdU as do controlled cells (Fig 6B).

**Fig 6:**
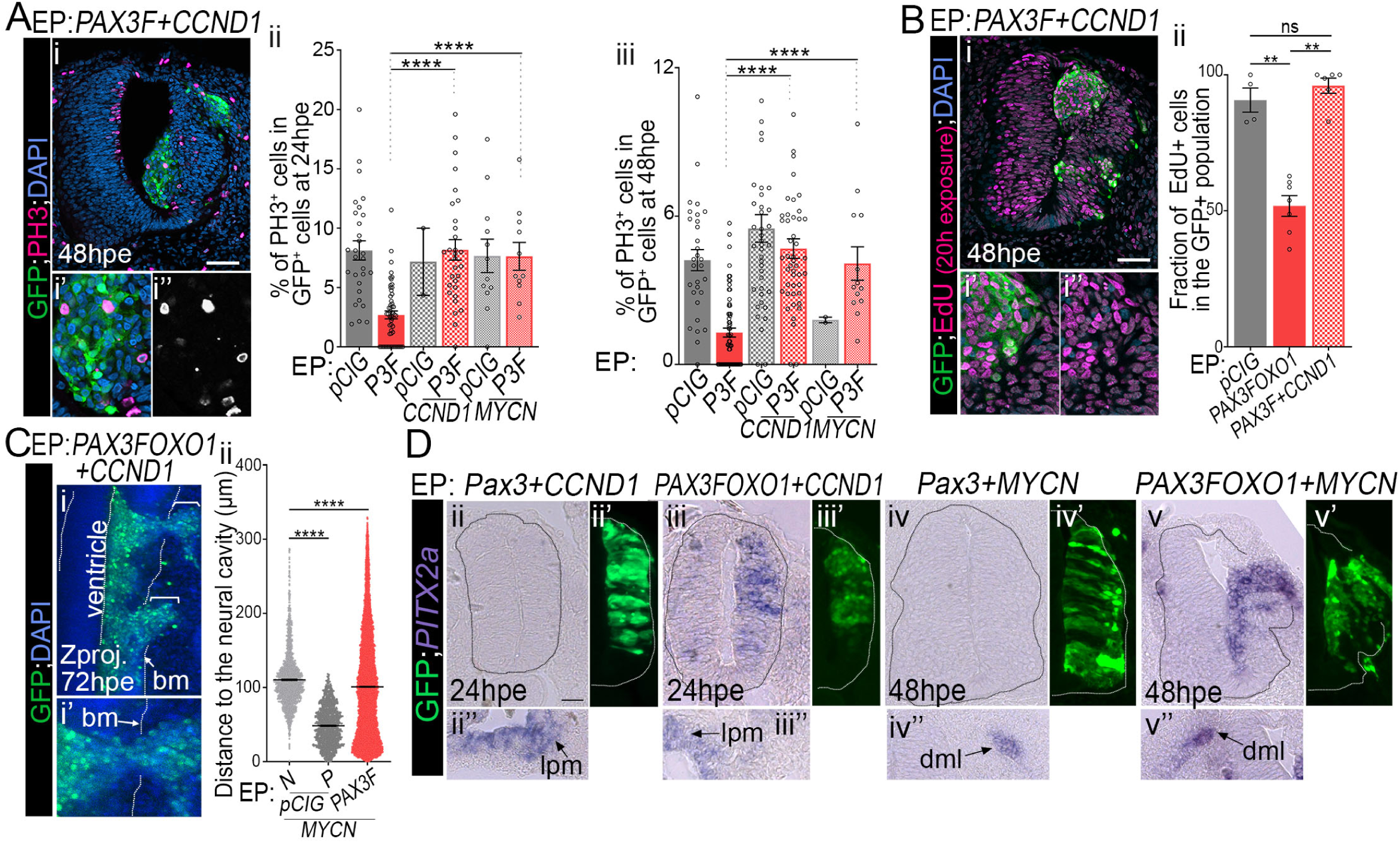
CYCLIN D1 and MYCN rescue PAXFOXO1 cell cycle inhibition, without affecting the identity and migration of cells. **(A) (i-i’’)** Immunodetection of GFP, the phosphorylated form of histone H3 (PH3) and DAPI staining on transverse section of chick embryo at 48hpe with *PAX3FOXO1* and *CCND1*. **(ii-iii)** Quantification of the number of PH3^+^ cells in the GFP^+^ cells in embryos expressing the indicated plasmids at 24hpe (**ii**) and 48hpe **(iii**). (dots: values on one section, n>8 embryos: mean ± s.e.m.). **(B) (i-i’’)** Immunodetection of GFP, EdU and DAPI staining on transverse section of chick embryos 48hpe with the indicated plasmids and soaked with EDU 20h before harvest. **(ii)** Quantification of the number of EdU^+^ cells in the GFP^+^ population in embryos expressing the indicated plasmids (dots: values on one section, n=3 embryos; histograms: mean ± s.e.m.). **(C) (i-i’)** Z projection along the dorso-ventral axis of 3D scans of an embryos 72hpe with *PAX3FOXO1* and *CCND1*. Dotted lines mark the apical cavity and the basal membrane (bm). **(ii)** Quantification of the distance of each GFP^+^ nuclei and the apical surface at 48hpe and 72hpe with the indicated plasmids (Violin plots). (**D**) *PITX2* detection via *in situ* hybridization on transverse sections of chick embryos 24hpe and 48hpe with the indicated plasmids and immuno-detection of GFP on the adjacent section slide. Bottom panels show areas on the embryos and section presented in the upper panels where PITX2 is expressed, including the lateral plate mesoderm (lpm) and the dorsal medial lip of the dermomyotome (dml). **Generals:** Scale bars 50µm; x’, x’’ panels are blown up on the box of the x upper ones. Mann-Whitney U test p-value: **: p< 0.01, ****: p< 0.0001, ns: p> 0.05.

We next wondered whether the proto-oncogenes recurrently amplified in ARMS cells could also overcome the G1 arrest of PAX3FOXO1 expressing cells. MYCN been amplified in about 10% of ARMS [6], we forced its expression together with PAX3FOXO1. In the neural tube, as previously demonstrated[60], MYCN, as opposed to its usual function, decreased the number of progenitors in M phase (Fig 6A). In contrast, in presence of MYCN, PAX3FOXO1^+^ cells became more actively proliferative (Fig 6A), with their rate of mitosis reaching levels comparable to that of control *pCIG* embryos.

Finally, we checked that upon reactivation of the proliferative activity of PAXFOXO1, the specific traits induced in the transformed neural progenitors were maintained. Assaying the expression of the ARMS marker gene *PITX2* and the migration of cells supported this idea (Fig 6C-D).

Together, these results indicated that PAXFOXO1 proteins inhibit the entry of cells into S phase and this is associated with a decrease in CDK-CYCLIN activity. This inhibition can be overcome by increasing the levels of CYCLINs or that of MYCN. This activity is not specific to the fusion proteins but can be also induced by a gain in the activity of the normal version of PAX3 and PAX7.

## Discussion

Characterising the early steps of cell response to PAXFOXO1s in chick embryos has highlighted the exceptional reprogramming potential of these factors. Within 72h, these factors induce the trans-differentiation of neural cells into ARMS-like cells that are capable of tissue remodelling and invasion. These activities may have been underappreciated in previous studies because of the focus on the long-term effects of PAXFOXO1s [13,29,30,32,33] and the negative impact of the fusion proteins on the cell cycle, a major brake in the formation of detectable primary tumour growths. Below, we discuss the relevance of the new model for understanding ARMS aetiology, as well as the molecular and cellular means by which the activity of these fusion TFs alters cell fate and behaviour.

### Embryonic neural cells respond to the ARMS transforming activity of PAXFOXO1

Our study provides evidence that embryonic spinal cord is responsive to PAXFOXO1 TFs activity. This in line with a report in zebrafish, where random insertion of *PAX3FOXO1* transgenes led to primitive neuroectodermal tumour formation, while the emergence of cell masses in skeletal muscle of the back required cooperating mutations [29]. Conversely, if the expression of PAX3FOXO1 from the murine *Pax3*^+^ spinal progenitors results in developmental defects, it is not sufficient for tumourigenesis [61,62]. The amount of PAX3FOXO1 produced by the activity of a single *Pax3* promoter, which is turned off when progenitors leave the cell cycle to become G0 neurons, are certainly too low to reach tumorigenic potential [28,29,32].

Strikingly, our data demonstrate that the high levels of PAXFOXO1s are able to switch on cohorts of TFs, whose expression is normally silenced in the neural tissue. This is likely stemming from a pioneer transcriptional activity [63], demonstrated for PAX3FOXO1 in human fibroblasts [12]. There, genomic recruitment of the fusion TF operates largely on closed and transcriptional shut down CRM. PAX3FOXO1 is able to displace the CRM nucleosomes and to set up an epigenetic landscape associated with active transcription. In agreement, the PAXFOXO1s were able to activate *de novo* CRMs in the embryonic neural tube.

In addition, our study supports the idea that the combination of genes induced by PAXFOXO1s is highly specific and reminiscent of that found in ARMS cells. Intriguingly, PAXFOXO1s activity can largely bypass the genomic constrain imposed by the origin of cell in which they act and can impose ARMS signature in various cell subtypes, including fibroblasts, mesenchymal stem cells or myoblasts, as well as various vertebrate species including chick, zebrafish, mouse and human [12,29,31,32](our data). This is further sustained by non-negligible overlap of PAX bound CRMs identified in discrete ARMS cell lines[12]. The originality of this ARMS signature stems from its unique association of determinants of several embryonic lineages, having in common a PAX3/7 dependent ontology [11,48](Fig 2C). The presence of PAX DNA motifs in at least half of PAX3FOXO1 bound CRMs represents a means by which PAX3/7 dependent development gene networks are co-opted by PAX3FOXO1 expressing cells [11,12]; a mechanism found recurrently during tumourigenesis [64]. This is further sustained by the formation of ARMS in the presence of fusion proteins made of the DNA binding domains of PAX3 and of either NCOA1/2 nuclear receptor co-activators or INO80D, a subunit of the INO80 chromatin remodelling complex [65].

Most importantly, our study, taken together with the study by Kendall *et al*. [29], supports the contention that neural tube cells are a cellular subtype from which ARMS can originate. Accordingly, 20-40% of primary tumour masses are found in organs colonized by NCC, such as the orbit, bladder, para-meningeal, head and neck areas ([40,46,66], S1A Fig). Moreover, several clinical studies report the presence of ARMS primary growths in a giant naevus and spinal cord, that are unambiguously neural tube derived [36,37]. This idea is further supported by the observation that the regulatory regions interacting with *PAX3* promoter upon t(2;13)(q35;q14) translocations can remain active in the neural tube after the translocation [67]. The impact of this cell origin on the manifestation of the disease and how much it can contribute the ARMS heterogeneity remain to be fully clarified, yet it is tempting to speculate that it will modulate tumour formation incidence, location and histology [29,33].

### PAXFOXO1s mediated cell cycle inhibition limits the expansion of transformed and metastatic cells

In the light of the cellular phenotypes appearing upon exposure to PAXFOXO1, we propose that these TFs stand as *sensus stricto* oncogenic drivers, whose activity is likely underpinning the time line of tumour formation. On the one hand, PAXFOXO1s rapidly provide cells with tissue remodelling and invasion capacities. This is reminiscent of the transformational power of a small group of TFs, named EMT-TFs [68]. Explaining this, PAXFOXO1 dependent ARMS signature is significantly enriched for key regulators of tissue remodelling. It includes notably modulators of RhoGTPases activity, such as ARHGAP25[47]. Indeed, the Rho GTPases are known to regulate cell-cell and cell-ECM interactions, polarity and migration [69], which are all modulated upon PAXFOXO1 exposure (Fig 4). In addition, PAXFOXO1 tissue remodelling activity could be reinforced by the recruitment of other EMT driving TFs, such as SNAI1, PRXX1, ETS1/2 [68] (Table S1, Fig 4). On the other hand, our analyses revealed that the oncogenicity of PAXFOXO1 transformed cells is limited by their low rate of proliferation (Fig 5). Such negative effect of PAXFOXO1s on cell cycle progression is unlikely to be inherent to our model system. Human myoblasts expressing PAX3FOXO1 are not to be able to produce colonies within soft agar [30], it takes several weeks of culture for PAX3FOXO1^+^ NIH3T3 cells to generate such colonies [70], and PAXFOXO1s^+^ human fibroblasts are arrested or spend more time in G1 phase, as do chick neural cells (Fig S3E). These results provide insight for why complementation of PAXFOXO1s with genetic aberrations promoting cell cycle progression, such as the gain of MYCN or CCND1, or loss of p53 or RB1, can enhance their tumorigenic potential (Fig 3, Fig 6) [28,30,34]. Whether such complementation is required for ARMS’ evolution and if so how it is achieved is not known for most cases. Alterations including mutations and small indels, copy number deletions and amplifications or structural variations within cell cycle regulators associated with PAXFOXO1 generating chromosome translocation is only seen in 30% of biopsies [5,6]. This calls for a better understanding of the molecular mechanisms underpinning this cell cycle inhibition. The buffered cell cycle progression induced by PAXFOXO1 proteins _a quasi dormant state_ could underlie the refractory response of ARMS cells to drugs such as CDK2 inhibitors [34] and contribute to the elevated resurgence of tumours post-treatment [71], as shown for other cancers [2]. We propose that RB1 activity inhibition is central to PAXFOXO1 mediated establishment of a dormant state. The decrease in the levels of the phosphorylated form of RB1 post PAX3FOXO1 gain of function points at a decrease in the level of CDK2 activity and explains the arrest or longer stay in early G0/G1 [72]. This is further supported by the elevated levels of CDKN1c (p57^Kip2)^, a protein that binds to and inhibits CDK2 activity [73], and was originally shown to cause cell cycle arrest mostly in G1 phase. This hypothesis is also compatible with a phenotypic rescue by a complementation with CCND1 (Cyclin D1), an efficient driver to S phase [74]. Strongly supporting the idea that RB1 regulation is a nodal point in PAXFOXO1 mediated cell cycle regulation, its loss of function have been shown to affect the progression, but not the formation of tumours from p53 null cells of the Myf6 embryonic muscle lineage overexpressing PAX3FOXO1[34].

Finally, amongst all the models that have been created to study ARMS development and evolution [28–30,32,33,75], our model uniquely recapitulates the invasive and disseminative properties of PAXFOXO1 expressing cells [71]. As previously demonstrated with human grafted cells [38], we believe that it will particularly suited for studying the modes of dissemination paths of PAXFOXO1 transformed cells. Our model is also foreseen to provide a means to dig into the molecular networks acting during the transition from a PAXFOXO1 mediated-latent metastatic state to overt metastasis [76]; and thereby to provide valuable insights for future therapeutics developments.

## Methods

### Bioinformatics

Transcriptomes of ARMS and ERMS biopsies have been published elsewhere [42–46] (accession numbers GSE92689, E-TABM-1202, E-MEXP-121 and data in [45]). Each dataset was based on Affymetrix micro-arrays. Details can be found at Table S1 (Sheet 2). Raw probe set signal intensities were normalized independently, using the frozen RMA method (fRMA Bioconductor R package [77]. Individual expression matrices were merged and the residual technical batch effects were corrected using the ComBat method implemented in the SVA R package [78]. Samples corresponding to tumour biopsies with validated presence/absence of *PAX3FOXO1* or *PAX7FOXO1* fusion genes were subset from the original data using a custom made R script. Differential analysis of fusion positive versus negative samples was conducted using the samr package and the following parameters: resp.type=“Two class unpaired”, nperms=100, random.seed = 37, testStatistic= “standard”; [79]. Genes with a delta score lower than 2.3 (FDR 0) where selected for subsequent analysis.

Hierarchical clustering of the normalized transcriptomes was implemented using the heatmap.2 function from the gplot package [80]). PAX3FOXO1 chIPseq data (GSE19063, Cao 2010) were mapped to human genome (hg19) using Bowtie2 [81] and peaks were called using MACS2 [82] implemented on Galaxy server [83]. Peaks common to the 2 replicates of Rh4 cell line and not present in the RD cell line samples were selected using BEDtools [84] and annotated to the two nearest genes using GREAT [85]. Functional annotation of the differentially expressed genes and the PAX3FOXO1 putative target genes was made using the analysis tool of the PANTHER Classification System [86] or GSEA [87].

### *Chick in ovo* electroporation

Electroporation constructs based on *pCIG* (*pCAGGS-IRES-NLS-GFP*) expression vector [88] have been described previously; *Pax3, Pax7, PAX3FOXO1, PAX7FOXO1* [61]; *MYCN* [60]; *CCND1* [89]. Reporters for the human or the mouse versions of PAX3FOXO1 bound enhancers were cloned upstream of the *thymidine kinase* (tk) promoter and *nuclear LacZ* [90] or *adenovirus major late promoter* (mlp) and *H2B-Turquoise*. For detailed cloning strategies see supporting methods. Reporter plasmids (0.5µg/µl) and *pCIG* based constructs (1.5-2 µg/µl) were electroporated in Hamburger and Hamilton (HH) stage 10-11 chick embryos according to described protocols [91]. Embryos were dissected at the indicated stage in cold PBS 1X.

### Immunohistochemistry and *in situ* hybridisation on cryosections

Embryos were fixed with 4% paraformaldehyde (PFA) for 45 min to 2 hr at 4°C, cryoprotected by equilibration in 15% sucrose, embedded in gelatin, cryosectioned (14 µm), and processed for immunostaining [91] or *in situ* hybridisation (ISH) [92]. Details of the reagents are provided in the Supp. Material and Methods. Immunofluorescence microscopy was carried out using a Leica TCS SP5 confocal microscope. Pictures of *in situ* hybridisation experiments were then taken with an Axio Observer Z1 microscope (Zeiss). All the images were processed with Image J v.1.43g image analysis software (NIH) and Photoshop 7.0 software (Adobe Systems, San Jose, CA, USA). All quantifications were performed using ImageJ v.1.43 g on usually more than 3 embryos and on 2 to 4 transverse sections per embryo. The number of cells positive for a marker per section in Fig 3Aiii, Biii, Fiii, S3Aiv,viii, Biv, Gi,ii was established using the cell counter plugin. Fluorescence intensities in GFP^+^ cells presented in Fig 1Evii, Fiii, 3Diii, Eiii, S1Fii,iv, S3Ci,ii, Div Fig were determined using ellipsoid region of interests whose size was adapted to that of cell nuclei and multi-measurement plugin. The levels of SOX2 and HUC/D in Fig 1Bv-vii were measured as described in [93] in bins of 2 cells wide and large of half of a neural tube placed in the dorsal region of the neural tube containing only GFP^+^ cells. All statistical analyses were performed using a Mann-Whitney U test in GraphPad Prism and all the p-values are given in figure legend.

### EdU pulse labelling and staining

A solution of EdU 500uM was injected within the neural tube lumen 20h before harvest. Immunofluorescence and EdU staining was performed as described previously [94] and with the Click-it EdU system (Thermo fisher).

### Cell dissociation from chick embryos

GFP positive neural tube regions were dissected after a DispaseI-DMEM/F-12 treatment (Stem cell technologies 1U/ml #07923; 37°C, 30min). Single cell suspensions were obtained by 3 minutes incubation in Trypsin-EDTA 0.05% (Life technologies) and mechanical pressure. Inhibition of Trypsin was ensured using with cold foetal bovine serum (FBS).

### RT-quantitative real-time-PCR on FAC sorted cells

GFP^+^ cells were sorted using BD Influx Sorter (BD Biosciences). Total RNA was extracted from 60 000 to 80 000 cells following RNAqueous-Micro kit with DNAseI (Life technologies) instructions. RNA quality was assessed by spectrophotometry (DeNovix DS-11 FX spectrometer). cDNA was synthesized by SuperScript VILO (Life Technologies) according to manufacturer’s instructions. RT–PCR was performed using the Veriti ™ 96-Well Fast Thermal Cycler (Applied Biosystems) and real-time qPCR was performed with the StepOnePlus™ real-time PCR system (Applied Biosystems) using SYBR Green detection tools (Applied Biosystems). Primers can be found in supporting methods. The expression of each gene was normalised to that of *TBP. ALK, CDC42, CDH3, CDK2, FOXF1, MYOD1, PITX2, RB1, TFAP2α, TFAP2β, TBP* expressions were assessed in n=6 (*pCIG*); n=4 (*Pax3*); n=6 (*PAX3FOXO1*) independent experiments. Other genes were tested in 3 independent experiments per condition. Data representation and statistical analyses using Mann-Whitney U-test or two-way ANOVA test were performed in GraphPad Prism.

### Flow cytometry-based cell cycle analysis

Dissociated cells were stained with 5uM Violet Vybrant™ DyeCycle™ (V35003, Thermo Fisher) at 37C for 30 min and Hoechst 33342. Fluorescent intensities and the related cell cycle phases determined were determined using a Cyan ADP Beckman Coulter analyzer. GFP negative cells served as an internal baseline for cell phase determination.

### GFP and DNA labelling and imaging 3D chick embryos

Samples were incubated overnight with Atto488 (1/300, Sigma) at 4°C for GFP staining, washed thoroughly in PBS, incubated 5-10 minutes in DRAQ5 (1/1000, Thermofisher) for DNA staining and finally washed in PBS. Samples were mounted on their ventral side in 1% agarose for 3D imaging. 3D scans of samples were obtained with a 2-photon microscope LaVision equipped with a femtosecond pulsed Insight Spectra Physics laser, a Carl Zeiss 20x, NA 1.0 (water immersion) objective and the InSight (LaVision BioTek) image acquisition software. A single wavelength of 930nm was used for exciting all fluorophores to avoid drift artefacts. Two GaAsp sensitive photomultipliers allowed simultaneous detection of the two emission lights which were segregated thanks to a dichroic mirror 585nm and a bandpass filter 525/50nm.

### 3D images processing and quantitative analyses

Image pre-processing and segmentation were performed using ImageJ and Imaris. Background subtraction was performed on GFP channel to eliminate autofluorescence coming from the tissue. Bleach correction normalizing the brightness of images along tissue thickness was performed on DRAQ5 channel stacks of thick samples, notably 72hpe samples. A quality filter on the Imaris automatic surface segmentation plugin based on intensity and size (<95 voxels) allowed removal of saturated unspecific objects and dead cells x,y,z coordinates of the centre point, the major axis of their ellipsoid fit, the sphericity and prolate ellipticity were retrieved for all segmented nuclei. The surfaces encompassing the neural tube, the neural cavity, pCIG progenitors and neurons were delineated on the DRAQ5 signal, on x-y planes every 3 z-stacks. Distance Transformation plugin outside neural cavity segmentation was used to quantify the distance between the centre of the nuclei and this cavity. *Cell clustering* was studied by running DBSCAN algorithm on Matlab [95]. Clusters contained a minima of 3 cells, and the minimal distance between two cells that belong to the same cluster was fixed to 10µm. *Cell orientation* was established by converting the cartesian coordinates of the vector representing the major axis of the ellipsoid fit of GFP positive cells into polar coordinates (S2C Fig) using Matlab. Matlab or Graphpad Prism were used for graphic representation and statistical analyses.

### Imaging the apical surface of Par3 and GFP labelled spinal cord

Dissected spinal cords were fixed in PFA4% for 1h and washed in PBS. Immunofluorescences were performed on cryosections. Open-book preparation of the samples flatten between a slide and coverslip was imaged using a spinning disk confocal microscope (Leica DMi8: CSU-W1 Yokogawa spinning disk) and MetaMorph (Molecular Devices) image acquisition software.

## Supporting information

SuppMat&Fig

## Acknowledgement

We deeply thank the ImagoSeine core facility of Institut Jacques Monod, a member of France-BioImaging (ANR-10-INBS-04) and certified IBiSA. We thank Griselda Wentzinger and Magali Fradet for performing cell sorting at ImagoSeine Institut Jacques Monod platform. We are grateful to the people who have provided us with useful tools. We received plasmids from Sophie Bel Vialar, Marie Henriksson, Elisa Marti, Gwen Le Dréau and YiPing Chen and ARMS and ERMS cell lines from Cécile Gauthier-Rouvière.

## Author contributions

**GGC:** Conceptualization; Formal Analysis; Investigation; Methodology; Software, Review & Editing, **ADV:** Conceptualization; Formal Analysis; Investigation; Methodology; Validation, Review & Editing, **YF:** Investigation; Formal Analysis, **LM:** Investigation; Formal Analysis; Software; Review & Editing, **LB:** Formal Analysis; Investigation; Methodology; Software, **SP:** Formal Analysis; Investigation; **NE:** Resources; Software, **FC:** Investigation, Methodology; Review & Editing, **MR:** Investigation, **FA:** Resources; Supervision, **ADR:** Resources; Supervision, **VC:** Methodology; Software, **FR:** Funding Acquisition, Editing, **OF:** Methodology; Supervision, **JB:** Conceptualization; Funding Acquisition, Review & Editing, **PGH:** Conceptualization; Writing – Review & Editing, **VR:** Conceptualization; Formal Analysis; Funding Acquisition; Investigation; Methodology; Project Administration; Supervision; Validation; Visualization; Writing – Original Draft Preparation.

