## Supplementary material for "The PAXFOXO1s trigger fast trans-differentiation of chick embryonic neural cells into alveolar rhabdomyosarcoma with tissue invasive properties limited by S phase entry inhibition": SuppMat&Fig

#### Supporting figures legends

**S1 Fig 1: (A) Body locations of RMS biopsies.** Locations of ARMS (red) and ERMS (blue) biopsies whose transcriptome has been assessed in Fig 2A and coming from previous studies [1–5]. **(B) PITX2 expression distinguishes ARMS from ERMS cells.** Pictures of western blots using the indicated antibodies on proteins extracted from the indicated ERMS (RD, RDAbl, Rh36) and ARMS (Rh3, Rh5, SJRH30, Rh4) cell lines, showing variable levels of PAX3FOXO1 (i) between ARMS cell lines and specific ectopic expression of several PITX2 isoforms (ii) in ARMS versus ERMS cell lines (see also S1 to S3_raw_images). **(C) PAX3FOXO1 activates TFAP2a and PAX2 expression in chick spinal cells (i,i',iii,iii')** Immunodetection of GFP, TFAP2α, and PAX2 on transverse sections of chick embryos 48hpe with *PAX3FOXO1*. **(ii,iv)** Quantification of expression levels of TFAP2α and PAX2 in $GFP^+$ cells in the spinal cords of chick embryos 48hpe with the indicated constructs (Dots: values in a cell, n>5 embryos; Mann-Whitney U test p-value: ****: p< 0.0001). Scale bar: 50µm.

**S2 Fig: Extended characterization of the epithelial-mesenchymal transition of PAXFOXO1 transformed embryonic spinal cells**

**(A)** Immunodetection of GFP and DAPI staining on transverse section of chick embryos 48hpe with the indicated plasmids. **(B) (i-iii)** Projection of 3D images of embryos 48hpe with the indicated plasmids, stained with DRAQ5 and immunolabelled for GFP. **(i'-iii')** Result of the segmentation performed at the level of the boxes indicated on samples i-iii. Surfaces delimiting the electroporated half of the neural tube are transparent yellow, while cell nuclei are coloured. In *pCIG* sample, the surface segregating progenitor nuclei from neurons is highlighted in transparent red. **(iv)** x (medial-lateral), y (antero-posterior) and z (dorsal-ventral) axes giving the orientation of i-iii samples. **(C) (i)** Representation in the 3 dimensions of the chick embryos of θ and φ polar angles of the vector (blue arrow) defining the major axis of a cell ellipsoid fit (blue circle). **(ii)** θ and φ possible values and major axes of chick embryos. **(iii)** Distribution of *φ* and θ in embryos electroporated with the indicated constructs at 12hpe and 48hpe, respectively. **(D)** Immunodetection for the indicated proteins at the indicated stages on transverse sections of chick embryos electroporated with the indicated plasmids. **(E)** Quantification of the distance of $GFP^+$ nuclei to the apical surface of the neural tube in embryos 72hpe with the indicated plasmids measured on transverse sections (dots: values in individual cells; histograms: mean ± s.e.m.; n>5 embryos). **(F)** Normalized levels of *SNAI1* and *ADAM10* mRNA assayed by DNA microarrays in ARMS and other type RMS biopsies (dots: value for a single RNA sample; histograms: mean ± s.e.m.). **Generals:** Mann-Whitney U test p-value: ****: p<0.0001, ns: p>0.05. Scale bars: 50µm; bottom x' and x'' panels are blown up of the boxes present on x panel.

**S3 Fig: Cell cycle state of PAX3/7 and PAXFOXO1 overexpressing embryonic spinal cells**
**(A) (ii-iv')** GFP and phosphorylated form of histone H3 (PH3) immunodetection and DAPI staining on transverse section of chick embryos at 48hpe (ii-iv') with the indicated plasmids. **(i, v)** Quantification of the number of $PH3^+$ cells in the $GFP^+$ population in embryos expressing the indicated plasmids at 24 (iv) and 48hpe (ii-iv'') (dots: value on a section; histograms: mean ± s.e.m.; n>8 embryos). **(B) (i-iii'')** GFP and activated CASPASE 3 immunodetection and DAPI staining on transverse section of chick embryos at 48hpe with the indicated plasmids. **(iv)** Quantification of the number of activated $CASPASE3^+$ cells in the $GFP^+$ population in embryos expressing the indicated plasmids at 48hpe (dots: value on a section; histograms: mean ± s.e.m.; n>8 embryos). **(C)** Quantification of the levels of the phosphorylated form of RB1 in the $GFP^+$ population in embryos expressing the indicated plasmids at 48hpe (dots: values in individual cells; histograms: mean ± s.e.m.; n>6 embryos). **(D) (i-iii'')** GFP and MCM2 immunodetection and DAPI staining on transverse sections of chick embryos at 48hpe with the indicated plasmids. **(iv)** Quantification of MCM2 levels in the $GFP^+$ cells after 48hpe with the indicated plasmids. (dots: values in individual cells; histograms: mean ± s.e.m.; n>5 embryos). **(E)** Proportion of cells in the indicated cell cycle phase assayed by DNA content distribution of FAC sorted $GFP^+$ chick neural (i-iii) and HFF cells (iv-vi) stained with the vibrant dye-cycle violet and the gating to separate cells in distinct cell cycle phase. Percentage of cells in the indicated cell cycle phase 48hpe established by FAC sorting GFP cells in embryos electroporated with the indicated constructs and stained with the Vybrant DyeCycle Violet$^{TM}$ (dots: mean value on cells analysed on independent FAC sorted samples; histograms: mean ± s.e.m.). **(F)** Heatmaps indicated the fold changes in the expression of the indicated genes relative to their mean expression in *pCIG* samples assayed in FAC sorted $GFP^+$ from chick embryos 48hpe with the indicated constructs. **Generals:** Mann-Whitney U test p-value: *: p< 0.05, **: p<0.01, ***: p<0.001, ****: p<0.0001, ns: p>0.05; Scale bars: 50µm; bottom x' and x'' panels are blown up of the boxes present on x panel.

### Supporting Methods

#### Cloning PAX3FOXO1 bound enhancers

Mouse versions of PAX3FOXO1 bound CRM nearby the *Met, Meox1, Myod1* and *Alk* genes were cloned upstream of the *thymidine kinase* (tk) promoter and *nuclear LacZ* and using the following primers:

*Met1$^{CRM}$-Fwd*: *TCCCAAGGCAGCTGCTACA*
*Met1$^{CRM}$-Rev*: *TGCGCTGTTTCCAGGGATC*
*Meox1$^{CRM}$-Fwd*: *CTCGAGGGAGTTGTTTCCT*
*Meox1$^{CRM}$-Rev*: *GCATGCTCCCGGCCGC*
*Myod1$^{CRM}$-Fwd*: *TCCAGAATGGGCTCGGTTC*
*Myod1$^{CRM}$-Rev*: *ACATGGTGACAAGGAATGGC*
*Alk$^{CRM}$-Fwd*: *TCTCCTTTTCAGCCACAGTG*
*Alk$^{CRM}$-Rev*: *TGGCCAGCAAGCTCCTT*

Human versions of PAX3FOXO1 bound CRM nearby the *CDH3* and *PRDM12* genes were cloned within the SacI and XhoI sites sitting upstream of *adenovirus major late promoter* (mlp) and *H2B-Turquoise* and using the following primers:

*PRDM12$^{CRM}$-FW: GCGAGCTCCTCCACTTCCCCTTCAATGT*
*PRDM12$^{CRM}$-Rev: CCGCTCGAGCGGCTCGTAGGACTTGAATA*
*CDH3$^{CRM}$-FW: GCGAGCTCCCGGCTAAGGGAATGCTC*
*CDH3$^{CRM}$-REV: CCGCTCGAGCGCTAAACATCATATCTGGCA*

#### Primer sequences used for RT-qPCR on chick FAC sorted neural cells

*Fw- ADAM10: CGATAATCCTGCTGTGCTCCTG*
*Rev- ADAM10: TGAAAGTCGAGGCGCAAGAAC*
*Fw- ALK: TGAGCAGTCTGGATCTCCCAA*
*Rev- ALK: TCCAGCTCACAAGGAGTCTCA*
*Fw-ARHGAP25: AGCCTGGAGTGCCTACTGAAA*
*Rev- ARHGAP5: CGTGCAAGTCCAAAGTCAGC*
*Fw-CCND1: TCCATCAGACCCGACGAGTT*
*Rev-CCND1: GGGGTCATTGCAGCCAGATT*
*Fw-CDK2: GTACAAGGCTCGCAACAAGC*
*Rev-CDK2: GAGCTTGTTCTCCGTGTGGA*
*Fw-EYA2: AGCCTGGAGTGCCTACTGAAA*
*Rev-EYA2: CGTGCAAGTCCAAAGTCAGC*
*Fw-FGFR4: GCGCAACTTCACCATCTCTGTA*
*Rev-FGFR4: AGCGTACAGCTTCTTGTCCAT*
*Fw-LMO4: ACCAAGAGCGGCATGATCC*
*Rev- LMO4: ATGATAGACATTGCCCTGTGCC*
*Fw-MEOX1: ACCTGACAAGGCTCAGGAGAT*
*Rev-MEOX1: TATGGCACTACTCGGAGCTGG*
*Fw- MYOD1: AGCCTGGAGTGCCTACTGAAA*
*Rev-MYOD1: CGTGCAAGTCCAAAGTCAGC*
*Fw-MYCN: TCTTCCCCTTCCCCGTCAA*
*Rev-MYCN: GAGCGTCTTTTCTCCACTGTCA*
*Fw-PAX2: ACCTGACGTGGTGAGACAAAG*
*Rev-PAX2: ACTACTTTGGGGGTCGCTACT*

*Fw-PITX2: AGATCGCCGTCTGGACCAA*
*Rev-PITX2: GGCTGCATCAGGCCATTGAA*
*Fw-PRDM12: CTGGTACGGGAACTCACACAAC*
*Rev-PRDM12: CACACGAAGGGCTTGTCCA*
*Fw-RB1: AACAGCGAGAGCCACGTAAA*
*Rev-RB1: TGCTTCTGCATTCTTGTTCGAG*
*Fw-SNAI1: TGTGTCTGCAAGATGTGCGG*
*Rev-SNAI1: GGAGCAGGTTTTGCACTGGT*
*Fw-TBP: TCGTGCCCGAAATGCTGAAT*
*Rev-TBP: TGCTCCTGTGCACACCATTT*
*Fw-TFAP2$\alpha$: CTCTGGAAGCTGACGGATAACA*
*Rev-TFAP2$\alpha$: GTCCTGAGACTGGGGGTAGAT*

#### Primer sequences used for qRT-PCR on ARMS cell lines

*Fw-ALK: TCTCATCGCAGCCGATATGG*
*Rev-ALK: GGCATCTCCTTAGAACGCTCT*
*Fw-ARHGAP25: CCTGGAGCACGGCCGGAATG*
*Rev-ARHGAP25: ACCACGGGCTCTGGGAGGTC*
*Fw-EYA2: ACCCCCAGTATTACGGCTCA*
*Rev-EYA2: TTTCGCTGGTGTGGAAGGTC*
*Fw-FGFR4: CCATAGGGACCCCTCGAATAG*
*Rev-FGFR4: CAGCGGAACTTGACGGTGT*
*Fw-FOXF1: CTCCCTGGAGCAGCCGTATC*
*Rev-FOXF1: ACTCCTTTCGGTCACACATGC*
*Fw-LMO4: GGCACGTCCTGTTACACCAA*
*Rev-LMO4: CGCCCTCATGACGAGTTCAC*
*Fw-MEOX1: GGGAGCACTGCCAATGAGAC*
*Rev-MEOX1: ATATCTGCGGAGCCGAGTCA*
*Fw-MYOD1: GAGCACTACAGCGGCGAC*
*Rev-MYOD1: TAGTAGGCGCCTTCGTAGCA*
*Fw-NHLH1: TTGAGCACCCAGAGGAGACT*
*Rev-NHLH1: CCACTTCAGGGTTCCATGGTC*
*Fw-PAX2: CTGGGGATTCCTCGCTCCAA*
*Rev-PAX2: CCACCTCCTCTAATGTGGGC*
*Fw-PAX3FOXO1: TCCAACCCCATGAACCCC*
*Rev-PAX3FOXO1: GCCATTTGGAAAACTGTGATCC*
*Fw-PITX2: GACTCCTTCGGAACTTGGCAC*
*Rev-PITX2: CCCAGAAGTAGCAGTTTGGCG*
*Fw-PRDM12: CACCGGAGCTGGATGACCTA*
*Rev-PRDM12: CCGTACCACACCAGCAGTTC*
*Fw-TBP: CACGAACCACGGCACTGATT*
*Rev-TBP: TTTTCTTGCTGCCAGTCTGGAC*
*Fw-TFAP2$\beta$: ATTTGAACCGGCAGCACACA*
*Rev-TFAP2 $\beta$: TGGGTCGGCTGTTCCCTATC*

##### Western Blots

ARMS and ERMS cells were lysed in a buffer composed of 10mM Tris-Cl pH7.5, 5mM EDTA, 150mM NaCl, 30mM Sodium pyrophosphate, 50mM Sodium fluoride, 10% glycerol, 1% NP40 and cOmplete$^{TM}$ Protease Inhibitor Cocktail (Sigma). Western blots were performed using the products and protocol by Thermo Fisher. They were revealed using the following primary antibodies: mouse anti-GAPDH (AB0067-20, Sicgen, 1:1000), mouse anti-Pitx2 (AF7388, R&D System, 1:1000), mouse anti-FOXO1 (CH-19, Cell signalling, 1:1000) and horseradish peroxydase-conjugated secondary antibodies (Jackson Immunoresearch).

##### List of antibodies used for immunostaining

| Antigen | Species | Dilution | Provider (reference) |
|---|---|---|---|
| ARL13b | Rabbit | 1/1000 | [6] |
| Cleaved CASPASE3 | Rabbit | 1/500 | Cell signalling (9661S) |
| CDH2 (N-Cadherin) | Mouse | 1/100 | Santa Cruz (sc-393933) |
| CDNK1c | Rabbit | 1/50 | Santa Cruz (sc-8298) |
| Activated β-CATENIN | Rabbit | 1/500 | Life Technologies (A-11132) |
| βGalactosidase | Mouse | 1/500 | Promega (Z378) |
| GFP | Chicken | 1/1000 | Abcam (ab13970) |
| Phospho- Histone H3 | Rabbit | 1/5000 | Millipore (06-570) |
| HUC/D | Mouse | 1/1000 | Thermo Fisher Scientific (16A11) |
| β1-INTEGRIN | Mouse | 1/1000 | Sigma (MAB19294) |
| LAMININ | Rabbit | 1/500 | Sigma (L9393) |
| LHX1/2 | Mouse | 1/20 | DSHB (4F2) |
| MCM2 | Rabbit | 1/500 | Ozyme (3619T) |
| PARD3 | Rabbit | 1/1000 | Millipore |
| PAX2 | Rabbit | 1/500 | Thermo Fisher Scientific (716000) |
| PAX6 | Mouse | 1/50 | DHSB (AB 528427) |
| Phosphorylated RB1 | Rabbit | 1/500 | R&D systems (MAB6495) |
| SOX2 | Rabbit | 1/200 | Thermo Fisher Scientific (48-1400) |
| TFAP2α | Rabbit | 1/100 | Santa Cruz (sc8975) |
| ***Secondary antibodies***: donkey against mouse, goat, guinea pig or rabbit IgG coupled to Alexa Fluorophores A488, A546, A568 or A647 (Thermo Fisher Scientific) were diluted 1:500 and used with DAPI (500ng/ml, Sigma, 28718-90-3). | | | |

##### *In situ* probes

Chick *PITX2a* isoform and *LMO4* probes were previously described [7,8].

#### Supporting Tables

**Supporting Table S1:** Gene expression levels in ARMS and ERMS biopsies and location of PAX3FOXO1 bound regions nearly the genes assayed.

***Sheet 1:*** Normalised expression levels of genes previously assayed using DNA-microarrays [1–5] in ARMS and ERMS biopsies

***Sheet 2:*** Origin of the samples and presence or not of PAX3FOXO1 or PAX7FOXO1.

***Sheet 3:*** Identity of the PAX3FOXO1 bound CRM (peaks) previously identified [9,10] nearby the genes assayed in Sheet1.

***Sheet 4:*** Position of PAX3FOXO1 bound CRM (peaks) on Hg19 genome.

**Supporting Table S2: Results of the Gene Ontology Biological Process term enrichment analysis.**

Sheet 1: Statistics for enriched terms related to cell identity, migration and cell cycle regulation.

Sheet 2: ARMS upregulated genes assigned to cell identity enriched terms.

Sheet 3: ARMS upregulated genes assigned to cell migration and adhesion enriched terms.

Sheet 4: ARMS upregulated genes assigned to cell cycle regulation enriched terms

### Supporting raw images

**S1_raw-image:** Full western blot membrane presented in S1B Fig_anti-FOXO1

**S2_raw-image:** Full western blot membrane presented in S1B Fig_anti-GAPDH

**S3_raw-image:** Full western blot membrane presented in S1B Fig_anti-PITX2

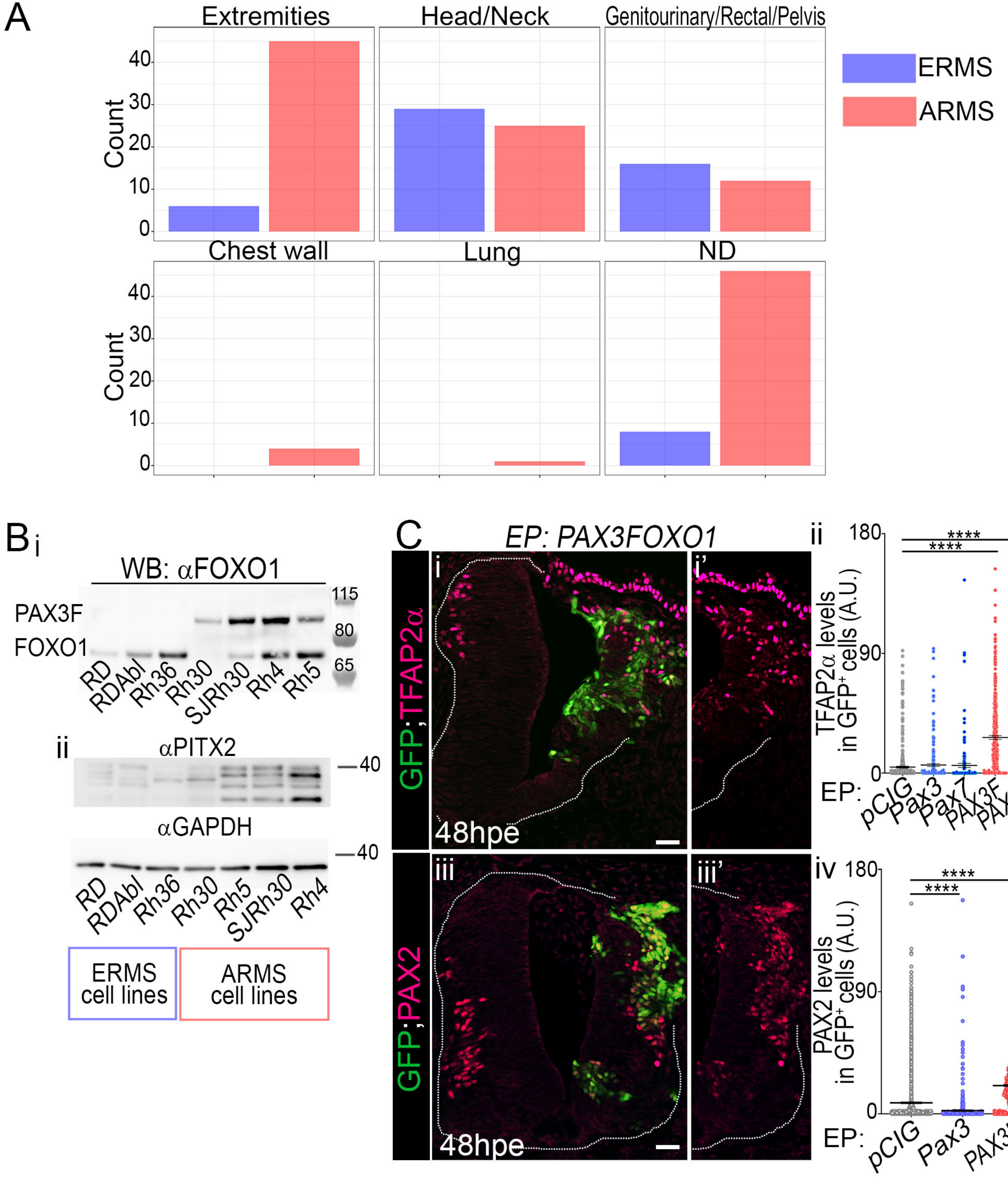


Curto et al., S1 Fig

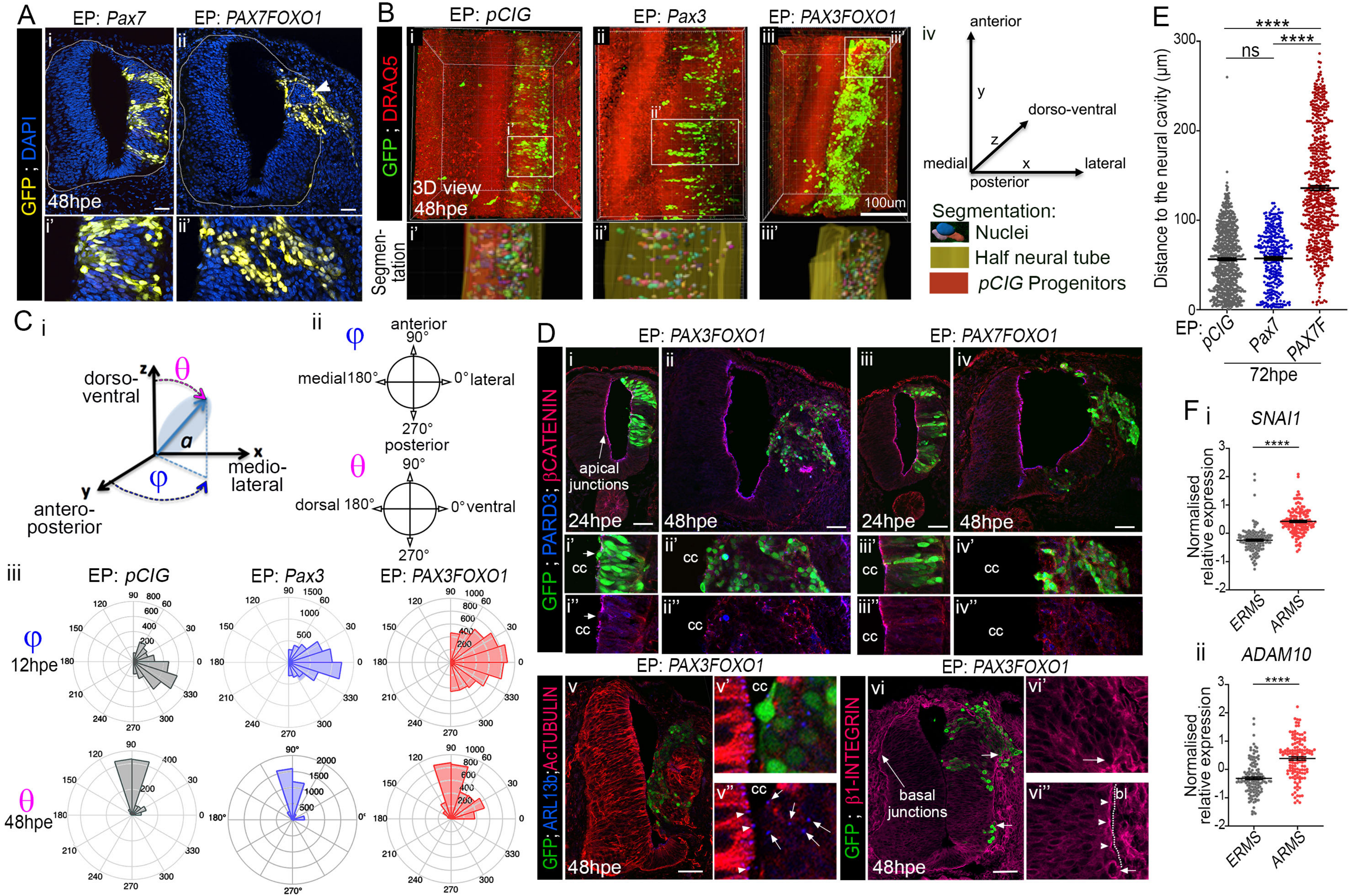


Curto et al., S2 Fig

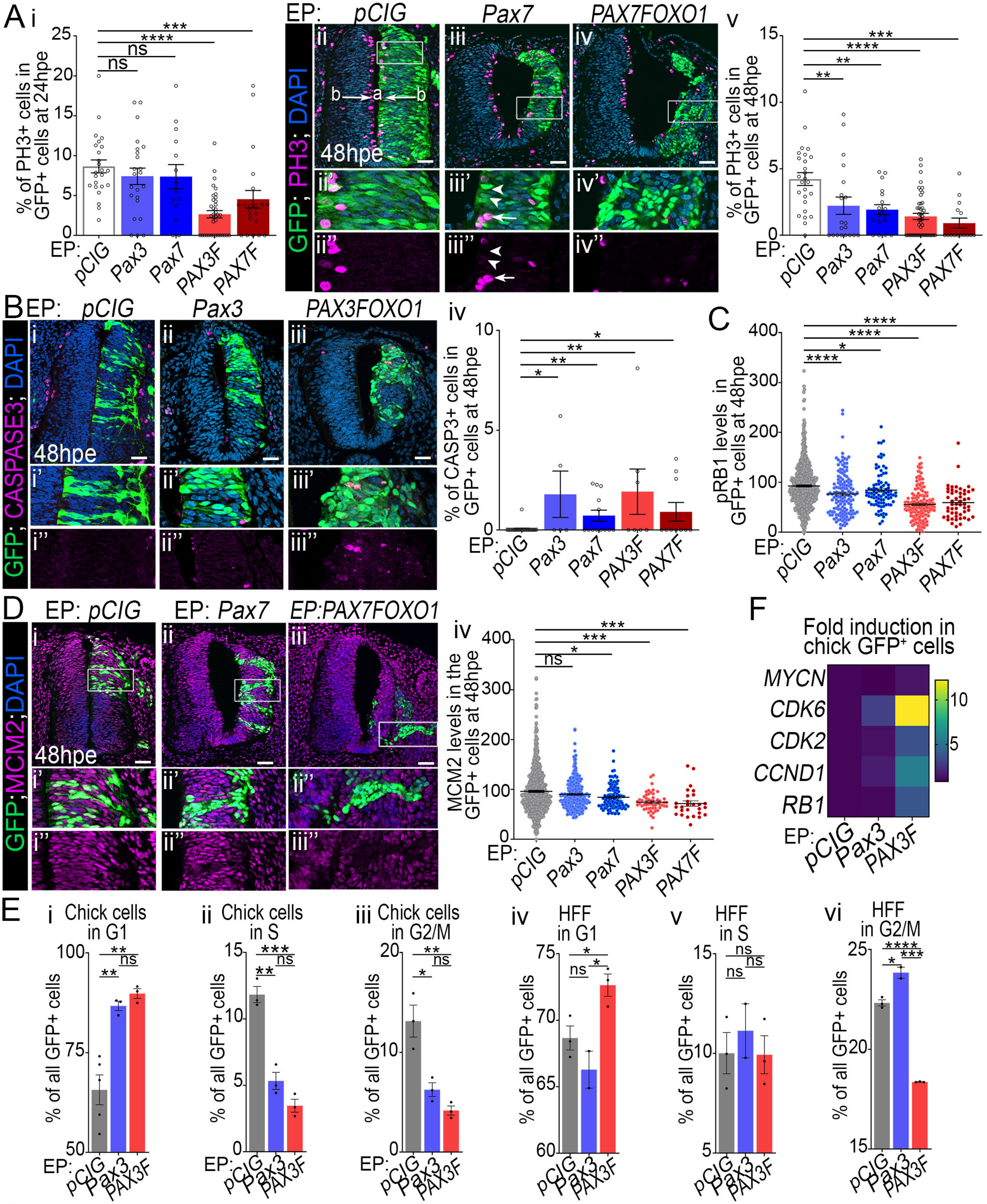


Curto et al., S3 Fig
